# Research Funding for Male Reproductive Health and Infertility in the UK and USA [2016 – 2019]

**DOI:** 10.1101/2021.08.23.456936

**Authors:** Eva Gumerova, Christopher J. De Jonge, Christopher L.R. Barratt

## Abstract

**Title:** Research Funding for Male Reproductive Health and Infertility in the UK and USA [2016 – 2019]

**Study question:** What is the research funding for male reproductive health and infertility in the UK and US between 2016 to 2019?

**Summary answer:** The average funding for a research project in male reproductive health and infertility was not significantly different to that for female-based projects (£653,733 in the UK and $779,707 in the US). However, only 0.07% and 0.83% of government funds from NIHR (UK) and NICHD (USA) was granted for male reproductive health, respectively.

**What is known already:** There is a marked paucity of data on research funding for male reproductive health.

**Study design, size, duration:** Examined government databases over a total 4-year period from January 2016 to December 2019.

**Participants/materials, setting, methods:** Information on the funding provided to male-based and female-based research was collected using public accessed web-databases from the UKRI-GTR, the NIHR’s Open Data Summary, and the US’ NIH RePORT. Funded projects that began research activity between January 2016 to December 2019 were recorded, along with their grant and project details. Strict inclusion-exclusion criteria were followed for both UK and US data with a primary research focus of male infertility, reproductive health and disorders, and contraception development. Funding support was divided into three research groups: male-based, female-based, and not-specified research. Between the 4-year period, the UK is divided into 5 funding periods, starting from 2015/16 to 2019/20, and the US is divided into 5 fiscal years, from 2016 to 2020.

**Main results and the role of chance:** Between January 2016 to December 2019, UK agencies awarded a total of £11,767,190 to 18 projects for male-based research and £29,850,945 to 40 projects for female-based research. There was no statistically significant difference in funding average between the two research groups (P=0.56, W=392). The US NIH funded 76 projects totaling $59,257,746 for male-based research and 99 projects totaling $83,272,898 for female-based research. There was no statistically significant difference in funding average between the two groups (P=0.83, W=3834).

**Limitations, reasons for caution:** The findings of this study cannot be used to generalize and reflect global funding trends towards infertility and reproductive health as the data collected followed a narrow funding timeframe from government agencies and only two countries. Other funding sources such as charities, industry and major philanthropic organizations were not evaluated.

**Wider implications of the findings:** This is the first study examining funding granted by main government research agencies from the UK and US for male reproductive health. This study should stimulate further discussion of the challenges of tackling male infertility and reproductive health disorders and formulate appropriate investment strategies.

**Study funding/competing interest(s):** CLRB is Editor for RBMO and has received lecturing fees from Merck, Pharmasure, and Ferring. His laboratory is funded by Bill and Melinda Gates Foundation, CSO, Genus. No other authors declare a conflict of interest.

## Introduction

Several recent studies have highlighted considerable research gaps in the understanding of male infertility encompassing critical areas such as basic science research, clinical diagnostics, non-Medically Assisted Reproduction (MAR) treatment options, and the impact of damage to the male genome on the health of the next generation (Barratt *et al*., 2017, Barratt *et al.*, 2018, De Jonge and Barratt 2019, Barratt *et al*., 2021, Schlegel *et al*., 2021a,b). One general conclusion that can be drawn from these analyses is that significant funding is required to address the research questions (Barratt *et al*., 2017, Barratt *et al.*, 2018). For any discipline, including reproductive medicine, an important aspect of assessing and formulating future funding requirements is to ascertain current funding. This knowledge can then be used to facilitate an objective needs analysis.

Surprisingly, there is a paucity of data on funding levels for male infertility and male reproductive health research (Barratt *et al.,* 2018, Barratt *et al.,* 2021). To date, only one study has specifically documented funding for male reproductive health research. Liao and colleagues (Liao *et al.,* 2020) assessed funding by the National Natural Science Foundation of China (NNSFC) for male infertility and reproductive health research between 1998-2018. The authors split this 20-year period into 3 funding phases beginning from 1998. By the third phase (2010-2018), a substantial increase of funding was awarded for male reproductive health (MRH) basic research by the NNSFC. However, there was minimal detail on the exact funding values. Barratt and colleagues provided a snapshot of funding for Male Reproductive Health in several countries that suggested overall funding levels were low, but no other details were provided (Barratt *et al*., 2021).

In this study, we investigated government funded support of male reproductive health research. We examined research funded between January 2016 to December 2019 from the UK and US agencies. To provide context, we included funding for female-based reproductive health research and examined the proportion of research funding for reproductive health research and compared to the total research funding.

## Materials and Methods

### Experimental Design

Publicly accessible UK Research and Innovation (UKRI), National Institute for Health Research (NIHR), and National Institutes of Health (NIH) funding agency databases covering awards from January 2016 to December 2019 were examined (see Supplementary Table I). Following the inclusion and exclusion criteria outlined within Supplementary Tables II and III, funding data were collected on research proposals investigating infertility and reproductive health. For simplicity, these are referred to collectively as ‘infertility research’. As the primary focus of this research is on infertility, the data were divided into three main groups: male-based, female-based, and not-specified (Supplementary Table II). The first two groups covered projects whose primary aim, based on the information presented in the research abstracts, timeline summaries and/or impact statements, was male- or female-focused. “Not-specified” includes research projects that have either not specified a primary focus towards either male or female or have explicitly stated a focus on both. The process was conducted and reviewed by E.G. with C.L.R.B. Total funding for all three groups, funding over time, and comparison with overall funding for a particular agency was examined.

### UK Data Collection

Starting in April 2018, the UK research councils, Innovate UK, and Research England were combined reporting under one organization, the UKRI (UKRI, 2019). The councils, such as the Medical Research Council (MRC), Biotechnology and Biological Sciences Research Council (BBSRC), Engineering and Physical Sciences Research Council (EPSRC), and Natural Environment Research Council (NERC), independently fund research projects according to their respective visions and missions; however, from 2018/19, their annual funding expenditures were reported under the UKRI’s annual reports and budgets. The UKRI’s Gateway to Research (UKRI-GTR) web-database allows users to analyse information provided on taxpayer-funded research. Relevant search terms such as “male infertility” or “female reproductive health” (see Supplementary Table II) were applied with appropriate database filters (Supplementary Table I). The project award relevance was determined by assessing the objectives in project abstracts, timeline summaries, and planned impacts. Supplementary Tables I, II and III provide the search filters and the reference criteria for inclusion/exclusion utilized for analysis. The UKRI-GTR provides the total funding amount granted to the projects within a designated period.

The Open Data Summary View dataset from the NIHR was used as it provided details on funded projects, grants, summary abstracts, and project dates. Like the UKRI data, the NIHR excel datasheet had specific search terms and filters applied to exclude irrelevant projects (Supplementary Tables I, II, and III).

The UKRI councils and NIHR report their annual expenditure and budgets for 1st April to 31st March. Thus, the selected projects will fall under the funding period of when their research activities begin, e.g. if a research project is started between May 20th, 2017, to March 20th, 2019, the project will be categorized under the funding period 2017/18. The projects assessed would begin their investigations between January 2016 to December 2019, therefore 5 consecutive funding periods were examined (2015/16, 2016/17, 2017/18, 2018/19, and 2019/20).

### US Data Collection

The NIH has a research portfolio online operating tools site (RePORT) providing access to their research activities, such as previously funded research, actively funded research projects, and information on NIH’s annual expenditures. The RePORT-Query database has similar features as the UKRI-GTR and NIHR such as providing information on project abstracts, research impact, start- and end-dates, funding grants, and type of research. The same inclusion-exclusion criteria were applied as for the UK data collection, (see Supplementary Tables I, II, and III).

In contrast to the UK funding agencies, the NIH’s fiscal year (FY) funding follows a calendar period from October 1st to September 30^th^, *i.e.,* FY2016 comprises funding activity from October 1st, 2015, to September 30th, 2016. Projects running over one calendar period are reported several times under consecutive fiscal years and the funds are divided according to the annual period of the project’s activity.

During data collection, 74 projects were found as active with incomplete funding sums as the NIH divides the grants according to the budgeting period of every FY. The NIH are in the process of granting funds for the FY2021, so projects ending in FY2020 or FY2021 have provided a complete funding sum. For the active projects ending after 2021, incomplete funding data is shown. It is assumed the funding will increase in value by the time the research project ends in the future. To remain consistent with the UK data, projects granted funding are totalled as one figure and recorded under the FY the project first began research, whether they are active or completed. Thus, the US funding is referred to as “Current Total Funding”. For the US, the initial data collection period ran between October 2020 to December 2020 but then restarted for a brief period in January 2021 to complete the remaining funding values for several of the active research projects.

### Data Analysis

The data were divided into the three groups and organized into the funding period or FY during which the project was first awarded. R-Studio (Version 1.3.1093) was utilized for the data analysis. Box-and-whisker plots are presented with rounded *P*-values. Kruskal-Wallis and Wilcoxon Rank Sum tests were generated to assess any statistical significance. The data were independently collected and do not assume a normal distribution, so rank-based, non-parametric tests such as the Kruskal-Wallis and Wilcoxon Rank-Sum were used. The Kruskal-Wallis test was used between more than 2 groups, with the *P-*values and Chi-Squared (χ^2^) values provided. The Wilcoxon test was used between two groups with the *P-*value and the Wilcoxon test statistic, *W,* included. *P*-values <0.05 were considered statistically significant.

## Results

### Total and Median Funding

#### UK Data

Total funding for infertility from the UK funding agencies and the summary statistics of the UK data are presented in Tables I and II. Table III details the proportion of funding by the MRC and NIHR from 2015/16 to 2018/19. Between 2016 to 2019, 76 studies were awarded funding by 4 UKRI councils and the NIHR investigating infertility and reproductive health. The MRC, BBSRC, and NIHR were the top 3 awarding agencies, having funded 29, 23, and 15 projects, respectively. The UK agencies have awarded 18 projects for male-based, 40 for female-based, and 18 projects for the non-specified group (Table I). For NIHR funding, there were only 2 awards for the male group compared to 11 for female group. Figure 1 presents a distribution of funding for the three groups. There was more spread for the female group, however there was no statistically significant difference between the mean values of the 3 groups (*P*=0.69, Kruskal-Wallis, χ^2^ = 0.72). There was no significant difference between male-based versus female-based funding (*P=*0.56, *W*=392).

**Table I.** Values collected are rounded to the nearest £ Sterling pound. [N] refers to the number of projects awarded. *Total funding for research grants and fellowships (in millions) by the respective UKRI councils and NIHR were determined by consulting their public annual budgeting reports; **As of the funding period 2018/19 and onwards, all UKRI councils report their annual expenditures as one, therefore the annual expenditure for research and innovation were obtained from the UKRI’s annual reports.

| Funding Period | Funding Agency | Total Funding for Research Grants and Fellowships (£000)* | Research Groups |  |  |
| --- | --- | --- | --- | --- | --- |
|  |  |  | Male-based (£)<br>[N] | Female-based (£)<br>[N] | Not-Specified (£)<br>[N] |
| 2015/16 | BBSRC | 330,473 | 0 | 0 | 0 |
|  | EPSRC | 691,280 | 0 | 0 | 0 |
|  | MRC | 487,157 | 1,748,922<br>[1] | 3,346,448<br>[1] | 0 |
|  | NIHR | 268,000 | 0 | 214,625<br>[1] | 0 |
| 2016/17 | BBSRC | 331,062 | 1,328,455<br>[3] | 0 | 695,839<br>[2] |
|  | EPSRC | 733,188 | 958,032<br>[1] | 0 | 0 |
|  | MRC | 341,630 | 676,826<br>[2] | 3,259,734<br>[6] | 1,444,459<br>[1] |
|  | NERC | 190,519 | 0 | 0 | 51,390<br>[1] |
|  | NIHR | 263,300 | 0 | 1,027,318<br>[1] | 0 |
| 2017/18 | BBSRC | 348,808 | 359,758<br>[1] | 359,772<br>[1] | 1,042,358<br>[2] |
|  | EPSRC | 844,134 | 694,461<br>[1] | 0 | 244,593<br>[1] |
|  | MRC | 325,164 | 903,026<br>[1] | 1,287,441<br>[2] | 636,510<br>[1] |
|  | NERC | 220,618 | 0 | 0 | 0 |
|  | NIHR | 274,000 | 477,541<br>[1] | 2,140,292<br>[3] | 0 |
| 2018/19 | BBSRC | 1,439,505** | 934,840<br>[2] | 1,719,275<br>[4] | 2,276,094<br>[4] |
|  | EPSRC/UKRI |  | 0 | 252,693<br>[1] | 0 |
|  | MRC |  | 0 | 5,535,724<br>[7] | 437,695<br>[1] |
|  | NIHR | 269,600 | 507,909<br>[1] | 165,595<br>[1] | 1,333,890<br>[2] |
| 2019/20 | BBSRC | 1,401,130** | 1,139,195<br>[2] | 779,165<br>[2] | 0 |
|  | EPSRC/UKRI |  | 1,230,976<br>[1] | 766,542<br>[1] | 1,455,327<br>[1] |
|  | MRC |  | 807,249<br>[1] | 1,564,917<br>[4] | 323,754<br>[1] |
|  | NERC |  | 0 | 0 | 611,514<br>[1] |
|  | NIHR | 321,200 | 0 | 7,431,405<br>[5] | 0 |
| **Total Funding (£)** |  |  | **11,767,190**<br>**[18]** | **29,850,945**<br>**[40]** | **10,553,423**<br>**[18]** |

**Table II:** All values are rounded to the nearest £ Sterling pound. SD, standard deviation of the data group; IQR, interquartile range which encompasses 50% of the data group.

|  | Research Group |  |  |
| --- | --- | --- | --- |
|  | Male-based<br>[N=18] | Female-based<br>[N=40] | Not-Specified<br>[N=18] |
| **Total (£)** | 11,767,190 | 29,850,945 | 10,553,423 |
| **Mean**<br>**(SD)** | 653,733<br>(384,131) | 746,274<br>(690,065) | 586,301<br>(387,951) |
| **Median**<br>**(IQR)** | 507,197<br>(423,630-789,793) | 476,163<br>(308,001-914,762) | 463,394<br>(328,457-630,261) |

**Table III:** The estimated proportion of funding for the UK was calculated using the data collected from Table 1. The proportions are rounded two 2 decimal points. The total research grants and fellowship values were obtained from the respective UK agency’s annual reports and budgets. The MRC total research grants and fellowships from 2018/19 – 2019/20 were excluded as they are part of the UKRI and report their annual expenditures under one with other research councils, therefore the exact sum for research grants and fellowships for MRC was not available. The total funding proportion is only looking at NIHR funding from 2015/16 to 2019/20.

| **Funding Period** | **UK Agency** | **Total Research Grants and Fellowships (£)** | **Total Funding Proportion for Infertility Research (%)** | **Male-based Funding Proportion (%)** | **Female-based Funding Proportion (%)** | **Not-Specified Funding Proportion (%)** |
| --- | --- | --- | --- | --- | --- | --- |
| **2015/16** | MRC | 487,157,000 | 1.05 | 0.36 | 0.69 | 0 |
|  | NIHR | 268,000,000 | 0.08 | 0 | 0.08 | 0 |
| **2016/17** | MRC | 341,630,000 | 1.58 | 0.2 | 0.95 | 0.42 |
|  | NIHR | 263,300,000 | 0.39 | 0 | 0.39 | 0 |
| **2017/18** | MRC | 325,164,000 | 0.87 | 0.28 | 0.4 | 0.19 |
|  | NIHR | 274,000,000 | 0.96 | 0.17 | 0.78 | 0 |
| **2018/19** | NIHR | 269,600,000 | 0.74 | 0.19 | 0.06 | 0.49 |
| **2019/20** | NIHR | 321,200,000 | 2.31 | 0 | 2.31 | 0 |
| **Total** | NIHR | 1,396,100,000 | 0.87 | 0.07 | 0.79 | 0.1 |

**Figure 1:**
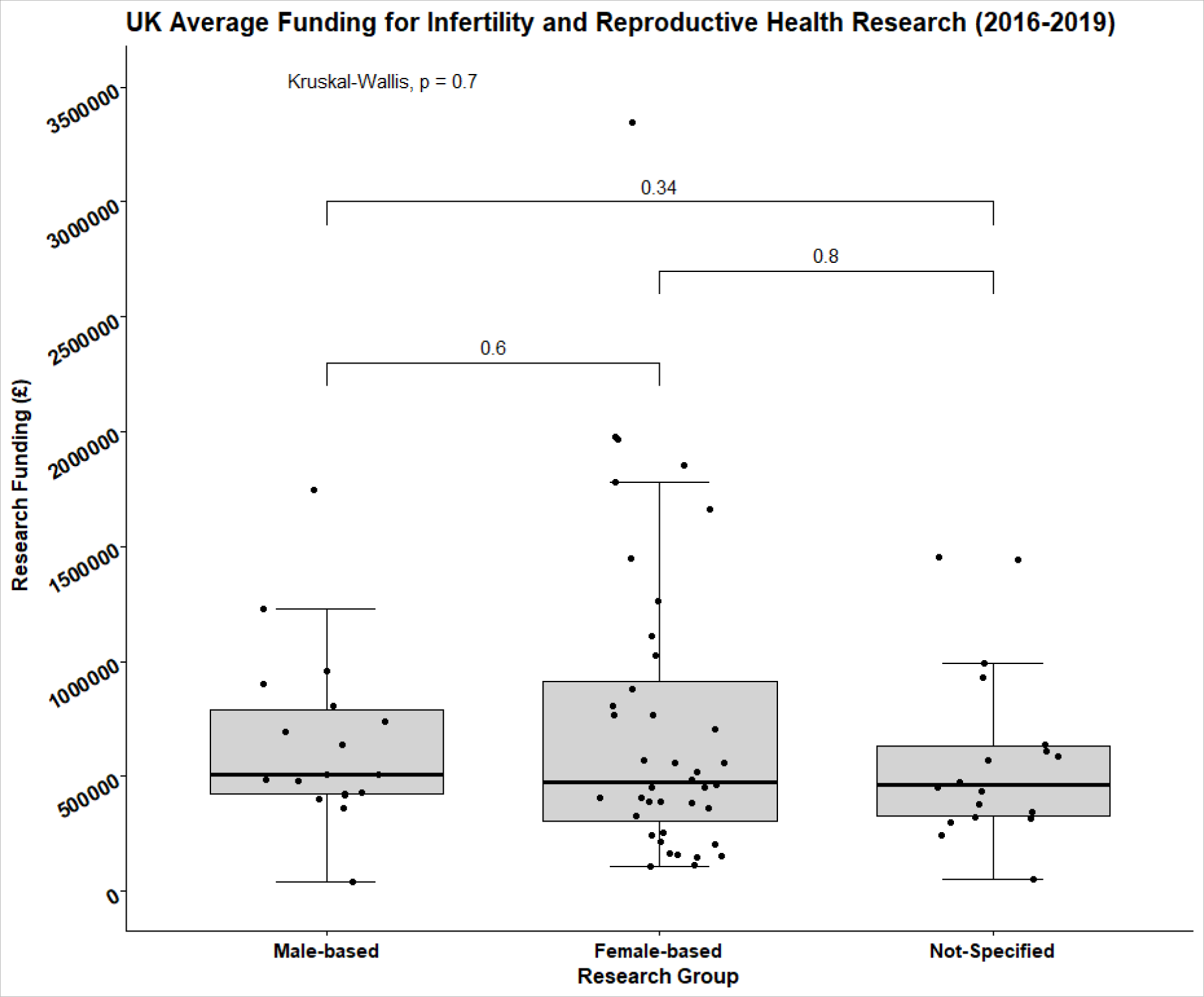
Box-and-whisker plot with a 95% confidence interval (CI) of awards for UK infertility and reproductive health research under the three research categories: male-based, female-based, and not-specified. 18 projects were funded for male-based research, 40 projects for female-based, and 18 for not-specified by the UKRI and NIHR.

#### USA Data

The US total funding for infertility and summary statistics are presented in Tables IV and V. The funding amounts presented in Table IV includes research grants, program grants, and fellowships and contains the respective annual spending of each NIH institute. The NIH have awarded 76 projects for male-based, 99 for female-based, and 31 projects for the non-specified group. The National Institute of Child Health and Human Development (NICHD), Environmental Health Sciences (NIEHS), and General Medical Sciences (NIGMS) have awarded the most for infertility research out of 14 institutes, funding 138, 27, and 26 projects, respectively.

**Table IV:**
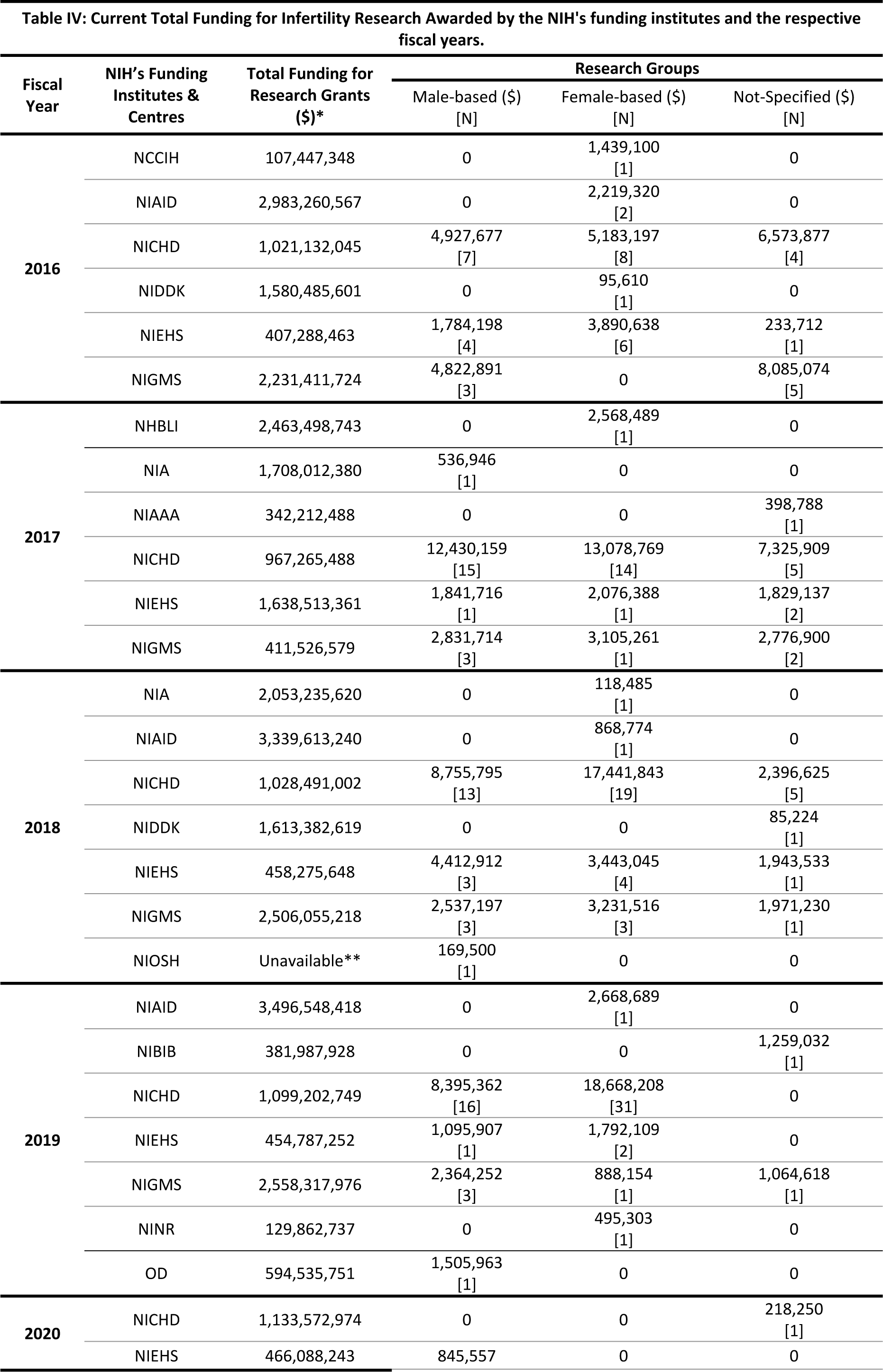
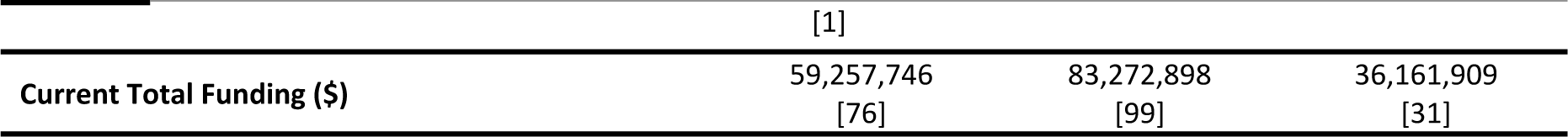
Values collected are rounded to the nearest US $ dollar. [N] refers to the number of projects awarded. From the start of data collection to the analysis, 7 projects changed their statuses from active to completed, making 138 projects out of 206 as active running. 67 of the 138 projects do not provide complete funding sums by the NIH, therefore, the funds were totalled up to their most recent awarding FY. *The values for the annual spending of research grants by the NIH (in millions) was found in the NIH’s RePORT Funding site: The Research Grants: Awards and Total Funding, by type and Institute/Centre. **The values were not made available by the NIH.

| Fiscal Year | NIH's Funding Institutes & Centres | Total Funding for Research Grants (\$)* | Research Groups | | |
| --- | --- | --- | --- | --- | --- |
| | | | Male-based (\$) [N] | Female-based (\$) [N] | Not-Specified (\$) [N] |
| 2016 | NCCIH | 107,447,348 | 0 | 1,439,100 [1] | 0 |
|  | NIAID | 2,983,260,567 | 0 | 2,219,320 [2] | 0 |
|  | NICHD | 1,021,132,045 | 4,927,677 [7] | 5,183,197 [8] | 6,573,877 [4] |
|  | NIDDK | 1,580,485,601 | 0 | 95,610 [1] | 0 |
|  | NIEHS | 407,288,463 | 1,784,198 [4] | 3,890,638 [6] | 233,712 [1] |
|  | NIGMS | 2,231,411,724 | 4,822,891 [3] | 0 | 8,085,074 [5] |
| 2017 | NHBLI | 2,463,498,743 | 0 | 2,568,489 [1] | 0 |
|  | NIA | 1,708,012,380 | 536,946 [1] | 0 | 0 |
|  | NIAAA | 342,212,488 | 0 | 0 | 398,788 [1] |
|  | NICHD | 967,265,488 | 12,430,159 [15] | 13,078,769 [14] | 7,325,909 [5] |
|  | NIEHS | 1,638,513,361 | 1,841,716 [1] | 2,076,388 [1] | 1,829,137 [2] |
|  | NIGMS | 411,526,579 | 2,831,714 [3] | 3,105,261 [1] | 2,776,900 [2] |
| 2018 | NIA | 2,053,235,620 | 0 | 118,485 [1] | 0 |
|  | NIAID | 3,339,613,240 | 0 | 868,774 [1] | 0 |
|  | NICHD | 1,028,491,002 | 8,755,795 [13] | 17,441,843 [19] | 2,396,625 [5] |
|  | NIDDK | 1,613,382,619 | 0 | 0 | 85,224 [1] |
|  | NIEHS | 458,275,648 | 4,412,912 [3] | 3,443,045 [4] | 1,943,533 [1] |
|  | NIGMS | 2,506,055,218 | 2,537,197 [3] | 3,231,516 [3] | 1,971,230 [1] |
|  | NIOSH | Unavailable** | 169,500 [1] | 0 | 0 |
| 2019 | NIAID | 3,496,548,418 | 0 | 2,668,689 [1] | 0 |
|  | NIBIB | 381,987,928 | 0 | 0 | 1,259,032 [1] |
|  | NICHD | 1,099,202,749 | 8,395,362 [16] | 18,668,208 [31] | 0 |
|  | NIEHS | 454,787,252 | 1,095,907 [1] | 1,792,109 [2] | 0 |
|  | NIGMS | 2,558,317,976 | 2,364,252 [3] | 888,154 [1] | 1,064,618 [1] |
|  | NINR | 129,862,737 | 0 | 495,303 [1] | 0 |
|  | OD | 594,535,751 | 1,505,963 [1] | 0 | 0 |
| 2020 | NICHD | 1,133,572,974 | 0 | 0 | 218,250 [1] |
|  | NIEHS | 466,088,243 | 845,557 | 0 | 0 |

Table IV: Values collected are rounded to the nearest US \$ dollar.
|  |  |  |  |
| --- | --- | --- | --- |
|  | [1] |  |  |
| Current Total Funding (\$) | 59,257,746 | 83,272,898 | 36,161,909 |
|  | [76] | [99] | [31] |

**Table V:** All values presented are rounded to the nearest US dollar and produced using RStudio.

|  | **Research Group** |  |  |
| --- | --- | --- | --- |
|  | Male-based<br>[N=76] | Female-based<br>[N=99] | Not-Specified<br>[N=31] |
| **Current Total (\$)** | 59,257,746 | 83,272,898 | 36,161,909 |
| **Mean** | 779,707 | 841,140 | 1,194,687 |
| **(SD)** | (594,203) | (862,707) | (1,046,679) |
| **Median** | 718,541 | 511,781 | 1,223,600 |
| **(IQR)** | (193,449-1,117,370) | (227,847-1,246,408) | (388,397-1,891,778) |

The spread of funding is not largely different between the male-based and female-based groups (Figure 2), but more projects appeared to localize at the lower end of the scale for the female-based group. However, there was no statistical difference between the mean values of the 3 groups (*P*=0.16, Kruskal Wallis χ^2^= 4.1). There were no significant differences between male- and female-based research *(P=*0.83, *W*=3834).

**Figure 2:**
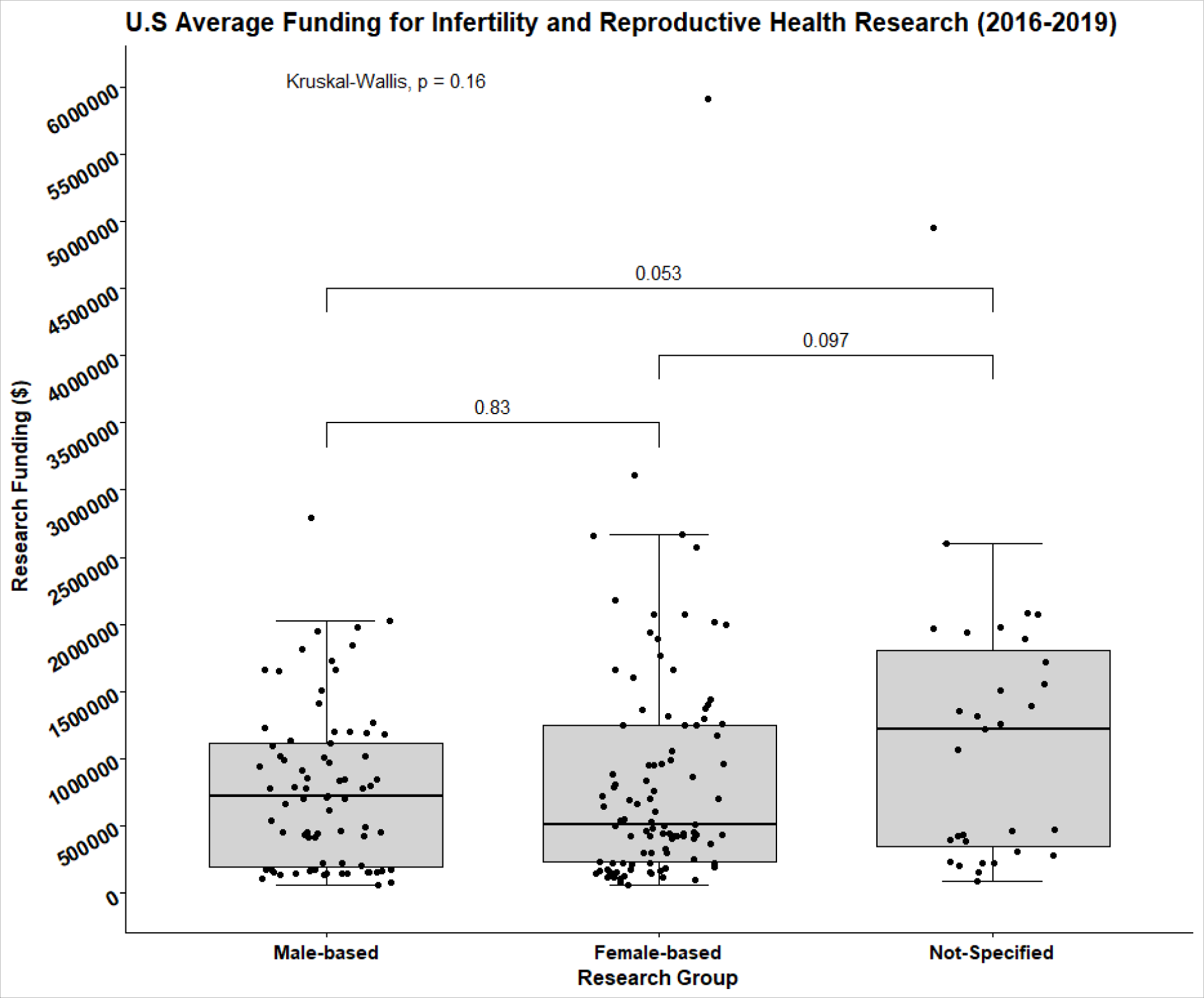
Box-and-whisker plot has a 95% CI of the funding collected for US infertility and reproductive health research under the three research focus categories: male-based, female-based, and not-specified. 76 projects were funded for male-based, 99 projects for female-based, and 31 for not-specified group by the NIH agencies.

### Funding over the years

Funding over 4 consecutive years is presented in Supplementary Tables IV and V for the UK and US, respectively. The total funding, mean funding amount over the respective funding periods, and the distribution of data are presented in Supplementary Figures 1 and 2. There were no statistical significant difference in the funding over time within each of the 3 groups *(P>*0.05, Kruskal-Wallis), for both the UK and US.

Proportion of Funding for Infertility and Reproductive Health Research in UK and USA:

The proportion of funding allocated to male and female infertility research is presented in Table III for UK and Table VI for US. The MRC fund research for reproduction and infertility and the NIHR have a dedicated research specialty for Reproductive Health and Childbirth (NIHR, 2021). When examining funding allocated directly for infertility research by the MRC, the proportion of total funding peaks at 1.58% in 2016/17 (Table III). For the NIHR, the largest proportion of funding allocated to infertility research was in 2019/20 with 2.31% of the year’s total awards. When examining total funding by the NIHR between 2015/16 and 2019/20, the proportion of funding for male-based infertility research was 0.07% and 0.79% for female-based research.

**Table VI:** The estimated proportion of funding for the US was calculated using the data collected from Table 4. FY2020 was excluded as only one project was awarded between 1^st^ October to 31^st^ December 2019 and would be unreflective of the funding proportion for this year. The total proportion looks at the total funding provided by the NICHD for infertility research from 2016 to 2019 and the total research funding granted by the NICHD. The annual research grants and fellowship values were obtained from the NIH’s RePORT Budget and Spending site: The Research Grants: *Awards and Total Funding, by type and Institute/Centre*.

| **Fiscal Year** | **NIH Agency** | **Total Research Grants and Fellowships (\$)** | **Total Funding Proportion for Infertility Research (%)** | **Male-based Funding Proportion (%)** | **Female-based Funding Proportion (%)** | **Not-Specified Funding Proportion (%)** |
| --- | --- | --- | --- | --- | --- | --- |
| **2016** | NICHD | 1,021,132,045 | 1.63 | 0.48 | 0.51 | 0.64 |
| **2017** | NICHD | 967,265,488 | 3.39 | 1.29 | 1.35 | 0.75 |
| **2018** | NICHD | 1,028,491,002 | 2.78 | 0.85 | 1.7 | 0.23 |
| **2019** | NICHD | 1,099,202,749 | 2.46 | 0.76 | 1.7 | 0 |
| **Total** | NICHD | 4,116,091,284 | 2.56 | 0.83 | 1.32 | 0.4 |

In the USA, of the 27 NIH institutes and research centres, the NICHD is a primary funder for furthering research on human development, improvement for reproductive health, and enhancing the lives of children and young adults (NIH, 2021). This also encompasses research for infertility and contraception development. The NICHD’s annual funding for research between fiscal years 2011 and 2020 was between $873 million to $1.1 billion (RePORT, 2021). In the FY2016, NICHD funded $1,021,132,045 for research grants and fellowships, but only 1.63% or $16,684,751 was for infertility research (as defined by the eligibility criteria in this study; Table VI). The funding proportion for the male-based research group was 0.48%, which was similar to the female-based funding proportion, 0.51%. The proportion of total funding provided by the NICHD between 2016 and 2019 that was allocated to infertility research was estimated at 2.56%, with male-based receiving 0.83% and female- based receiving 1.32%.

## Discussion

This study provides details of UK and US government funding for male infertility and male reproductive health covering the period 2016-2019. The information will be instructive for different stakeholders, *e.g*. workers in the discipline, grant organisations, commercial companies, and policy makers. This will enable the development of evidence-based informed decisions for future funding strategies This is critical as male infertility poses a global health risk for many millions of men yet research funding is not concomitant with the prevalence or impact of the disease.

We analysed public-accessible databases for UKRI, NIHR (UK) and US (NIH) covering the period of awards from January 2016 to December 2019. The primary objective was to determine funding for male reproductive health and infertility research. To provide context, we assessed 3 groups based on the primary focus of the research in reproductive biology/medicine: male-based, female-based, and not-specified (Supplementary Table II). Information from the aims, research abstracts, timeline summaries, and/or impact statements, was used to determine if a study was included and, if so, to which group it was assigned. This is necessarily a subjective process, therefore we provide our search and entry/exclusion criteria (Supplementary Tables I, II, and III), as well as a supplementary table of the research projects’ titles from the UK and US (Supplementary Tables VI and VII). Whilst incorporation of different terms may produce different answers, the results are robust. For example, the application of data extraction is consistent between countries as the inclusion/exclusion criteria were the same. We were focused on infertility and associated links to infertility and reproductive disorders. No analysis was made to assess if there is bias in funding research for female reproduction versus male reproduction. Moreover, we do not examine submission numbers, triage, rejection rates, etc. and thus prioritization of research cannot be assessed.

Although the number of awards for female-based research is generally higher than for the male group (ratio of ∼2:1 in UK, and 1.3:1 in USA), the average amount awarded per project was not significantly different in either country (see Tables II and IV; Figures 1 and 2), indicating that funding per project is not different between male and female reproductive health.

An important question to answer is, what is the proportion of funding for reproduction/male reproductive health compared to general research funding? There are several approaches to address this question. For both the UK and USA data, one method is to examine the total funding for research by the main funding agency and compare this to the data for male- and female-based research. Reproductive health research is primarily supported by the MRC and NIHR in the UK, and by the NICHD in the USA. In the funding periods 2015/16 to 2017/18, the total infertility research funding by MRC ranged from 0.87% to 1.58% of the total budget (Table III). Infertility research funding from NIHR ranged from 0.08% to 2.31% (2015/16 to 2019/20, Table III). For the US, the maximum infertility research funding by the NICHD was 3.39% of its total budget (Table VI).

Another approach was to assess the proportion of funding compared between research disciplines, or research categories, in the UK and USA, respectively. Within the UK data, we specifically examined research disciplines funded by the NIHR. From the April 1^st^, 2011 to present, the NIHR awarded over £216 million for Reproductive Health and Childbirth research, their 7th largest funding category. Mental Health, Cancer, and Cardiovascular Diseases were within the top 5 most funded categories (Supplementary Table VIII). NIHR awarded £21 million in 2017/18 for Reproductive Health and Childbirth research (NIHR, 2021), yet surprisingly there was minimal support towards male-based research as between 2016 to 2019 only two projects were funded (Table I, Supplementary Table VI). The small number of projects in male reproductive health funded by the NIHR was unexpected as NIHR are the largest UK funders for health care and clinical research (NIHR, 2021). NIHR supported 302 studies for reproductive health with 94 of them being newly funded projects for 2019/20. However, using our criteria for study inclusion we only identified 4 projects focusing on infertility over the whole period (Table I, Supplementary Table IV). While we do not know the reason for the low funding rate, a plausible explanation is that, as NIHR fund a significant number of clinical trials, there may not have been sufficient high quality candidates for either diagnostic and/or treatment trials to be developed in male reproductive health (Barratt *et al*., 2021).

To compare different research categories for the USA data, we did not use our collected data to provide estimated funding. Instead, we used the NIH’s *Research Portfolio Online Reporting Tools* estimates of funding for various Research Condition and Disease Categories (RCDC) (https://report.nih.gov/funding/categorical-spending<u>#/</u>) and the NIH’s annual research grants (https://report.nih.gov/funding/nih-budget-and-spending-data-past-fiscal-years/budget-and-spending). For the NIH, the values presented for the 299 RCDCs are not mutually exclusive because a project can fall under several categories. We examined research categories similar to those at the NIHR. For NIH these included: Contraception/Reproduction, Infertility, Obesity, and Mental Health (Supplementary Table IX). By estimating the proportion of funding for these categories from the NIH’s Total Research Funding, we can see those categories such as Obesity and Mental Health were highly funded in comparison to Contraception/Reproduction and Infertility.

NICHD has funded under 1% of their annual research grants for male-based research for 3 out of 4 consecutive fiscal years (Table VI). NICHD are primary funders for reproduction, infertility, and contraceptive development, therefore, it was unexpected to observe such low funding proportions. A possible factor for why our calculated funding proportion values by the NICHD are low may be due to our strict eligibility criteria during data collection. However, we applied our eligibility and exclusion criteria equally across all funding agencies, for the UK and US.

Two pertinent points arise from our study. Firstly, compared to the prevalence of the disease where 1:7 couples are infertile (Boivin *et al*., 2007, NICE, 2013), the proportion of research funding for male reproductive health is small (less than 1%, see Tables III and VI) compared to other diseases in the UK and US (Supplementary Table IX). This is surprising especially because MAR is a multi-billion-dollar global industry. Secondly, although the number of awards for female-based research is generally higher than for the male group (ratio of ∼2:1 in the UK and 1.3:1 in the USA), the average funding awarded per project is not significantly different in either UK or USA (see Tables II and IV; Figures 1 and 2). Whilst there are many challenges in comparing research funding between disciplines, the present findings directly imply a significant gap between impact of disease prevalence and research funding to investigate the disease, e.g., diagnosis and treatment. This apparent gap requires further detailed analysis and should include a comprehensive assessments of the health economic impact of male reproductive health.

There are several limitations to our study. Firstly, these findings cannot be generalized to reflect funding trends towards infertility and reproductive health worldwide. The data were collected from governmental agencies of two countries and over a narrow funding period. Further, the funding priorities of UK and US governmental agencies may not be a ‘good fit model’ for the funding priorities of government research agencies in other countries. Secondly, only government funding was investigated. We did not examine funding from non-governmental organizations (NGO’s), *e.g*. Wellcome Trust, industry, Bill and Melinda Gates Foundation, and other major philanthropic organisations. As the UKRI, NIHR, and NIH are governmental agencies, their prioritization to providing fellowships, research grants, program centre grants, and others may not be similar to other charities and international organizations. Detailed analysis of funding from these other agencies would be instructive and assist in a more comprehensive analysis. Future work should include data from more countries, NGO’s and include longer funding timeframes to accurately estimate total funding supporting for male infertility and male reproductive health and for more comprehensive assessment of funding trends.

In summary, we present recent government funding for male-based infertility and reproductive health, and by extension, funding towards female-based research. The information provided in this study should be useful for a variety of stakeholders as discussed earlier. A sentinel message is that whilst male infertility poses a global health risk for many millions of men, research funding to develop better diagnostic tools and treatment regimens is not on par. The data analysis presented herein should stimulate discussions for a strategic development of male reproductive health care investments.

## Acknowledgements

The initial concept for this work was based on discussions as part of the ESHRE MRHI Working Group. We are grateful to ESHRE for providing ground-breaking funding and for their continued support. We are grateful to Dr Stephen J. Publicover of University of Birmingham for providing critical feedback on the manuscript.

## Authors Roles

The experimental design, primary data collection, and initial statistical analysis was performed by EG as part of her undergraduate BSc honours research project. The initial draft of the manuscript was written by EG and CB following discussions with CJD. All authors contributed to writing and editing the manuscript and approving the final version.

## Data Availability Statements

The data underlying this article are available in the Dryad Digital Repository at https://doi.org/10.5061/dryad.v9s4mw6wc.

## Abbreviations List

**Abbreviations List:**
| **Abbreviations for Funding Agencies/Institutes, Research Councils, and Databases** |  |
| --- | --- |
| BBSRC | Biotechnology and Biological Sciences Research Council |
| EPSRC | Engineering and Physical Sciences Research Council |
| HRCS | Health Research Classification System |
| MRC | Medical Research Council |
| NCCIH | National Centre for Complementary and Integrative Health |
| NERC | Natural Environment Research Council |
| NIA | National Institute of Aging |
| NIAAA | National Institute on Alcohol Abuse and Alcoholism |
| NIAD | National Institute of Allergy and Infectious Diseases |
| NIBIB | National Institute of Biomedical Imaging and Bioengineering |
| NICHD | National Institute of Child Health and Health Development |
| NIDDK | National Institute of Diabetes and Digestive and Kidney Diseases |
| NIEHS | National Institutes of Environmental Health Sciences |
| NIGMS | National Institute of General Medical Sciences |
| NHBLI | National Heart, Blood, Lung Institute |
| NIH | National Institutes of Health |
| NIHR | National Institute of Health and Research |
| NINR | National Institute of Nursing Research |
| NIOSH | National Institute for Occupational Safety and Health |
| OD | NIH’s Offices of the Director |
| RCDC | Research, Conditions and Diseases Categories |
| RePORT | Research Portfolio Online Reporting Tools |
| UKRI | UK Research and Innovation |
| UKRI-GTR | UK Research and Innovation – Gateway to Research |

## Supplementary Materials

**Supplementary Table I.** N/A – not applicable to the database used or not an available search filter. *Training, institutional is the support of research training programs within the research areas and priorities supported by the Institute to train predoctoral and/or postdoctoral fellows of an institution. ** ‘Limit Project search to’ is a feature of the NIH RePORTER Query system that searches the inputted search terms at the following options: project title, terms, and abstracts.

| **Supplementary Table I: Search filters used for respective databases** |  |  |  |
| --- | --- | --- | --- |
| **Search Filters** | **UKRI – GTR** | **NIHR – Open Data** | **NIH – RePORT Query** |
| **Project Start Year/Date** | 2016;<br>2017;<br>2018;<br>2019 | 2016;<br>2017;<br>2018;<br>2019 | >=01/01/2016 |
| **Research Type**<br>**Or Program Type** | Research Grants;<br>Fellowships | Research;<br>Training | Research Projects<br>Grants;<br>Training, Individual;<br>Training, Institutional* |
| **Limit Project search**<br>**to\*\*** | N/A | N/A | Project Title<br>Project Terms<br>Project Abstracts |
| **Fiscal Year (FY)** | N/A | N/A | No FY was selected |
| **Funder**<br>**Or**<br>**Agency/Institute/Centre** | None Chosen | N/A | Check All |

**Supplementary Table II.** The inclusion criteria for the 3-research groups after examining abstracts, research impacts and public health relevance statements. Under each heading are the topics or field of research of reproductive health and infertility research.

| **Male-focused infertility research** | **Female-focused infertility research** | **Not-specified infertility research** |
| --- | --- | --- |
| Spermatogenesis research | Oogenesis research | Gametogenesis and infertility |
| Sperm maturation, motility, and quality analysis research | Menstruation/menstrual cycle abnormalities | EDCs impacting general fertility |
| Sperm dysfunction/disorders research | Uterine and Ovarian disorders (*e.g., Endometriosis, Polycystic Ovarian Syndrome, and Uterine Fibroids*) | Underpinning the aetiology of infertility (*not specifying the focus group*) |
| Testicular and Prostate health (not cancer) research | Female contraception research development | Contraception development in conjunction with HIV/STI prevention |
| Male contraception research development | Maternal factors associated with ART outcomes | Studies using animal and insect models to further the understanding of **human** infertility and contraception development |
| Paternal factors associated with ART outcomes | EDCs association with female infertility | Hormonal dysregulation |
| Paternal association with fetus health/development (i.e., male infertility affecting fetus health/development) | Underpinning the aetiology of female infertility | Cell/embryonic development dysregulation giving rise to infertility |
| EDCs giving rise to male infertility | Hormonal dysregulation and female infertility |  |
| Underpinning the aetiology of male infertility |  |  |
| Hormonal dysregulation and male infertility |  |  |

**Supplementary Table III.**
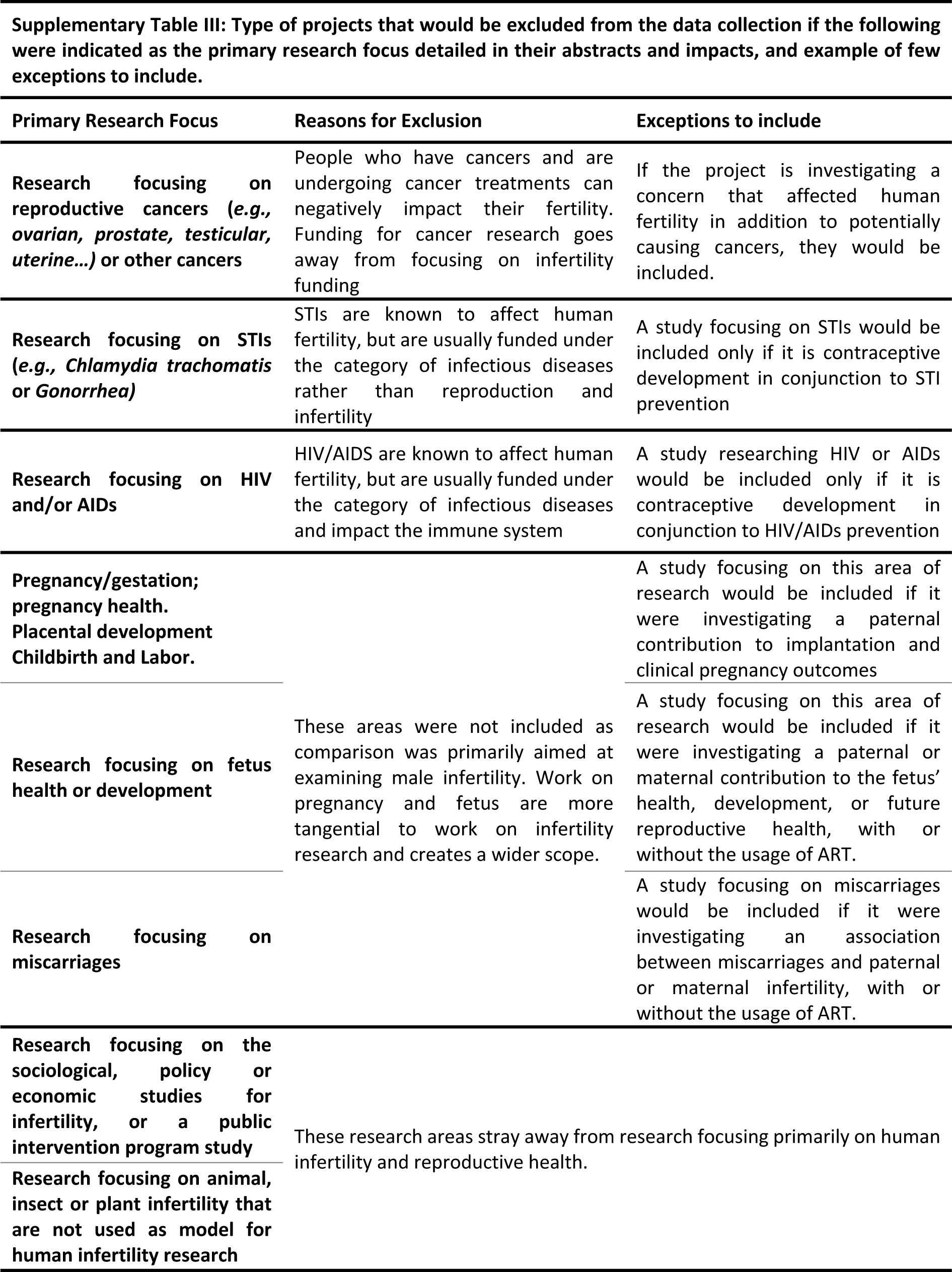
Type of projects that would be excluded from the data collection if the following were indicated as the primary research focus detailed in their abstracts and impacts, and few exceptions to include them.

**Supplementary Table IV.** The UK funding across the 5 consecutive funding periods awarded between January 2016 to December 2019. The values are rounded to the nearest Sterling pound. *Value belongs to a single project awarded, thus, no mean was produced.

| Funding Period | Research Groups |  |  |
| --- | --- | --- | --- |
|  | Male-based (£)<br>[mean] | Female-based (£)<br>[mean] | Not-Specified (£)<br>[mean] |
| **2015/16** | 1,748,922* | 3,561,073<br>[1,780,534] | 0 |
| **2016/17** | 2,963,313<br>[522,539] | 4,287,052<br>[562,710] | 2,191,688<br>[547,922] |
| **2017/18** | 2,434,786<br>[580,108] | 3,787,505<br>[631,251] | 1,923,461<br>[480,865] |
| **2018/19** | 1,442,749<br>[480,917] | 7,673,287<br>[590,253] | 4,047,679<br>[578,240] |
| **2019/20** | 3,177,420<br>[794,355] | 10,542,029<br>[878,502] | 2,390,595<br>[796,865] |
| **Total** | 11,767,190 | 29,850,945 | 10,553,423 |

**Supplementary Table V.** The US current funding for projects awarded funding beginning between January 2016 to December 2019. The values are rounded to the nearest US dollar. *For 2020, male-based and not-specified groups were each awarded 1 project since the beginning of the FY (October 1^st^ to December 31^st^, 2019), thus, no mean or median was produced.

| **Fiscal Year** | **Research Groups** |  |  |
| --- | --- | --- | --- |
| | Male-based (\$)<br>[mean] | Female-based (\$)<br>[mean] | Not-Specified (\$)<br>[mean] |
| **2016** | 11,534,766<br>[823,912] | 12,827,865<br>[712,659] | 14,892,663<br>[1,489,266] |
| **2017** | 17,640,535<br>[882,027] | 20,828,907<br>[1,225,230] | 12,330,734<br>[1,233,073] |
| **2018** | 15,875,404<br>[793,770] | 25,103,663<br>[896,559] | 6,396,612<br>[799,574] |
| **2019** | 13,361,484<br>[636,261] | 24,512,463<br>[680,902] | 2,323,650<br>[1,161,825] |
| **2020\*** | 845,557 | 0 | 218,250 |
| **Current Total** | 59,257,746 | 83,272,898 | 36,161,909 |

**Supplementary Table VI.**
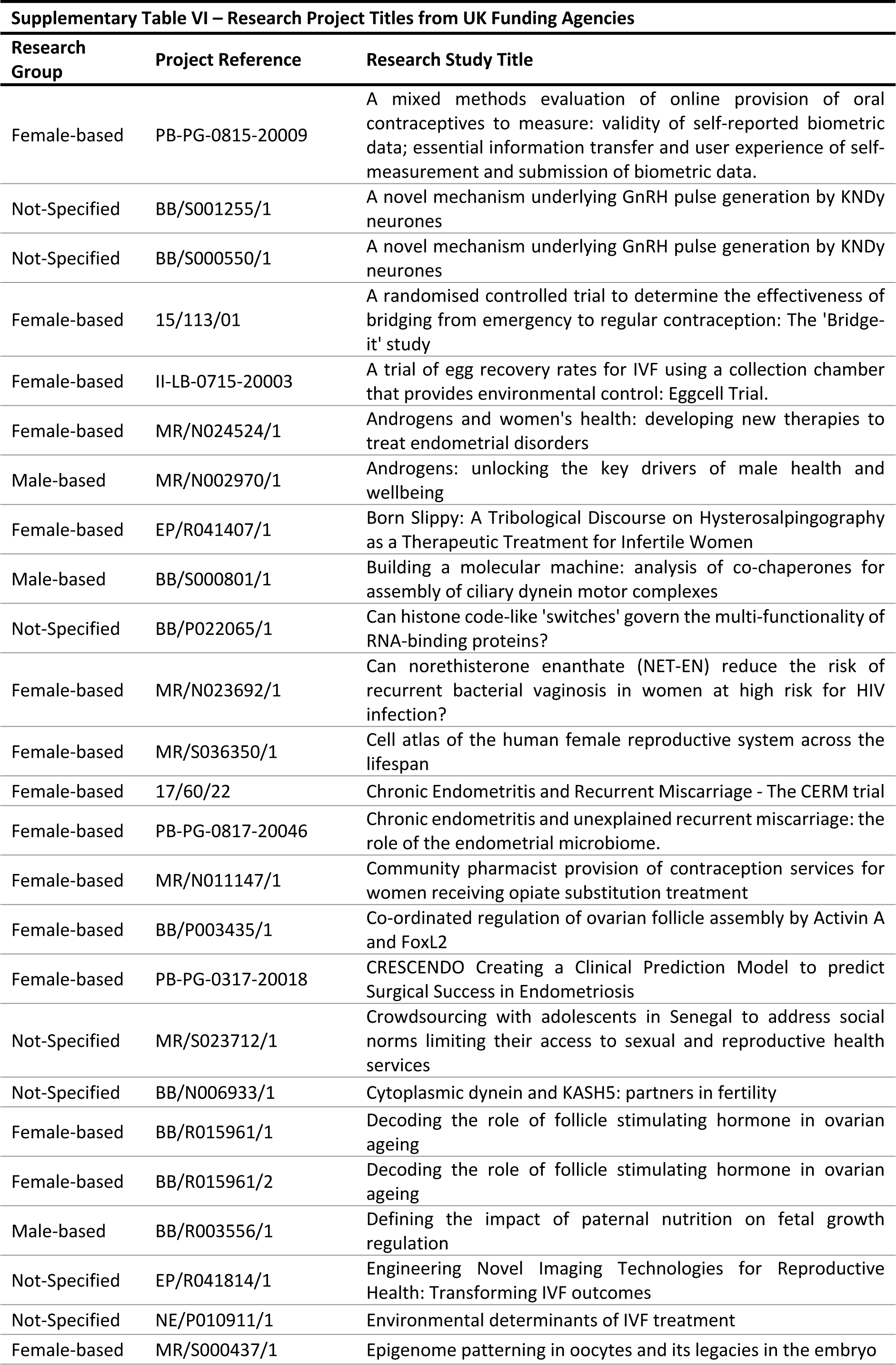
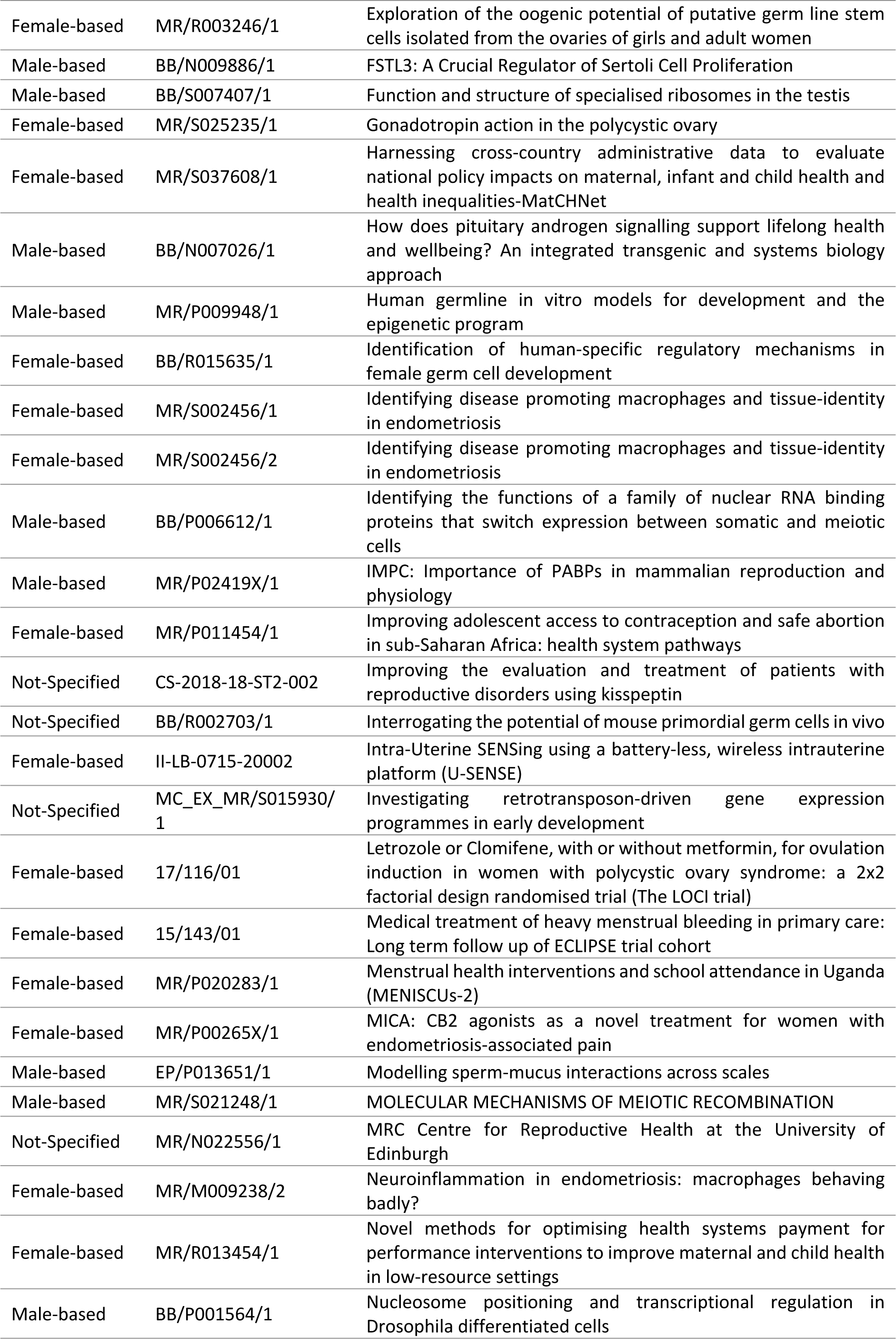
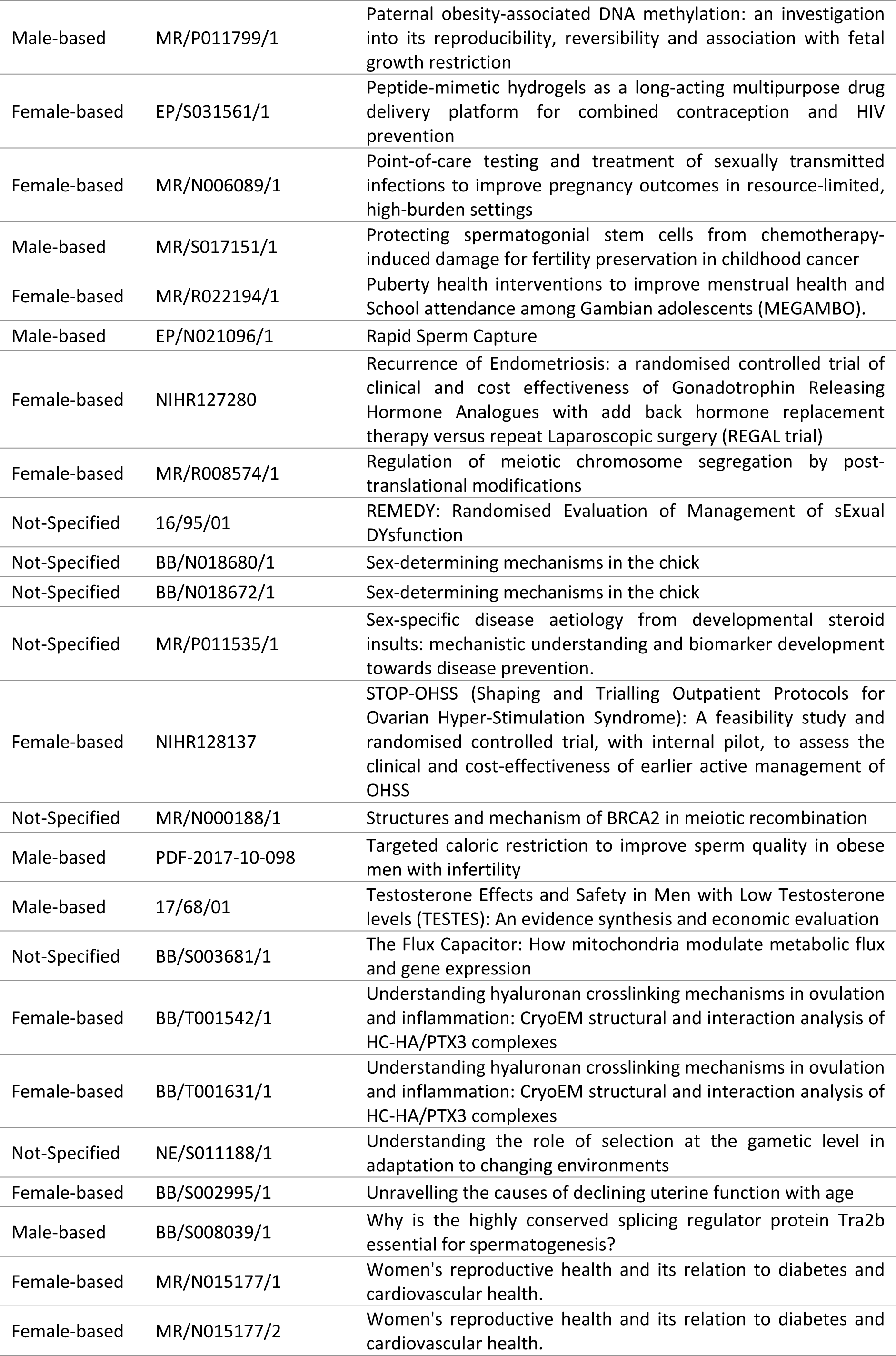
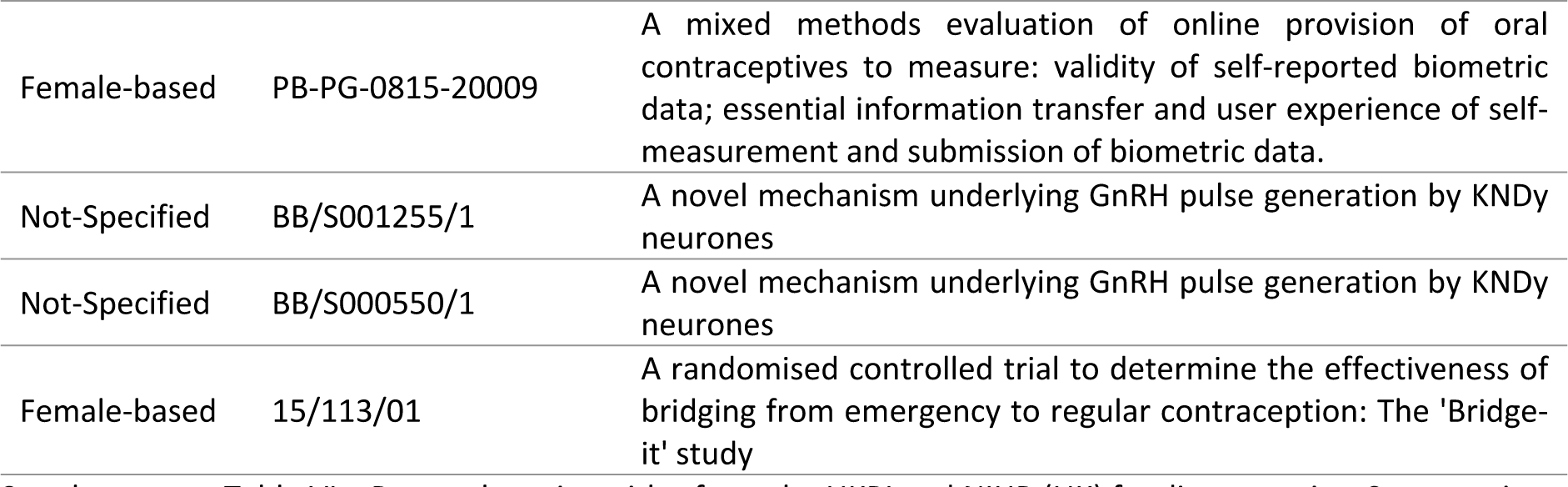
Research project titles from the UKRI and NIHR (UK) funding agencies. Some projects titles are repeated; however they would have separate grants awarded with different project IDs. Further information on each research project can be found in the UKRI and NIHR Dataset_EG_CDJ_CLRB.

**Supplementary Table VII.** Research project titles from the NIH (US) funding agencies. Some projects titles are repeated; however they would have separate grants awarded under different project IDs. Further information on each research project can be found in NIH Dataset_EG_CDJ_CLRB.

| Research Group | Project Reference | Research Study Title |
| --- | --- | --- |
| Female-based | 1R01HD100318-01 | 3/4: Pre-IVF Treatment with GNRH Antagonist in Women with Endometriosis - A prospective double blind placebo controlled trial (PREGNANT) |
| Male-based | 1R03HD087429-01A1 | A genome-wide drosophila RNA1 screen for regulators of centrosome reduction |
| Male-based | 1R21HD092828-01 | A home-based, rapid, sensitive semen analyser |
| Female-based | 1R01ES026964-01 | A longitudinal study of endocrine disrupter mixtures and reproductive aging |
| Female-based | 1R21HD090242-01 | A mouse model to demonstrate the impact of myometrial FSHR on fertility |
| Female-based | 1R01ES028923-01A1 | A preconception cohort study of air pollution, fertility, and miscarriage |
| Not-Specified | 1F32HD095620-01 | A role for hypothalamic melanocortin 3 receptors in integrating energy state with reproductive physiology |
| Male-based | 5R01HD091068-03 | A system for culturing mammalian spermatogonial cells |
| Male-based | 1R01HD095841-01 | Accumulation, storage, and release of sperm in the oviduct |
| Male-based | 1R01HD098039-01 | ALDH1A/A2 inhibitors for male contraception |
| Male-based | 1R61HD099743-01 | Allosteric cdk2 inhibitor discovery and development for male contraception |
| Female-based | 1F32HD090854-01A1 | An ovary-on-a-chip for understanding early folliculogenesis and reproductive toxicology in a large mammalian model |
| Female-based | 1U54HD096957-01 | Antibody-based contraceptive MPTS: preclinical and clinical research |
| Male-based | 1F31HD089693-01A1 | Assessing microenvironment and endothelial cell instruction of testis function |
| Female-based | 1R01HD096266-01 | Biological role of uterine glands in pregnancy |
| Not-Specified | 275201500002I-4-27500002-1 | Biological testing facility - maintenance of existing non-human primate colony |
| Female-based | 1R01AT008824-01A1 | Botanicals derived progestins and their impact on women's health |
| Male-based | 1R01HD096745-01A1 | Calcium signaling nanodomains in sperm motility and fertility |
| Not-Specified | 1R44HD097063-01 | Capsule-intravaginal ring for sustained release of antibodies for non-hormonal contraception and vaginal protection against HIV |
| Not-Specified | 1R35GM118066-01 | Causes and consequences of aneuploidy |
| Female-based | 1R01HD092550-01 | Causes and consequences of mitochondrial dysfunction in oocytes and cumulus cells |
| Female-based | 275201300019I-0-27500005-1 | CCTN-pharmacokinetic/pharmacodynamic evaluation of levonorgestrel butanoate for female contraception |
| Male-based | 1R15HD096759-01A1 | Cell-type specific inactivation of sumoylation during mouse spermatogenesis |
| Male-based | 1F32HD086986-01 | Chromatin dynamics, transcriptional activators and repressors in transition from proliferating progenitors to terminal differentiation during adult stem cell differentiation |
| Female-based | 1R01GM123048-01A1 | Chromatin expulsion by the DNA replication licensing factor orc4 during asymmetric cell division in meiosis and differentiation |
| Male-based | 1R01GM123643-01 | Circumventing the blood-testis barrier |
| Female-based | 1F30HD085652-01A1 | Compartmental adrenomedullin signalling in the uterus during implantation |
| Male-based | 1R21HD090371-01A1 | Comprehensive mapping of mouse testis cell types and spermatogenic stages by single-cell rna sequencing |
| Male-based | 1R01GM127379-01A1 | Control of spermatogonial stem cell formation |
| Female-based | 1R61HD099750-01 | Copper intravaginal contraception |
| Not-Specified | 1R21HD098621-01A1 | Correction of endocrine disruptor-induced transgenerational epimutations by CRISPR-DCAS9 |
| Male-based | 1P01HD087157-01A1<br>8120 | Crispr/cas9 and Small Molecules for Targeting Sperm Function and Fertilization |
| Male-based | 1R01GM123512-01A1 | Deciphering Pachytene piRNA Function |
| Male-based | 1R03HD093990-01A1 | Defining Mechanisms of Pp1 Phosphatase Specificity and Function Required for Male Fertility |
| Female-based | 1F31HD097830-01 | Defining the Neurophysiologic Mechanisms Engaged by Estradiol Feedback in Regulating Reproductive Neuroendocrine Function |
| Not-Specified | 1R15HD084253-01A1 | Defining the Novel Role for the Rna Binding Protein Etr-1 in C.elegans Gametogenesis |
| Male-based | 1R01OD028223-01 | Derivation of Functional Spermatogonia Stem Cells From Rhesus Macaque Ipscs |
| Male-based | 1R01ES030942-01 | Determining How Preconception Exposure to Phthalates Impacts Sperm Function, the |
| Female-based | 1R21ES026454-01A1 | Detrimental Effects on Female Reproduction of in Utero and Neonatal Exposure to Common Phthalates DEHP and Its Replacement DINP |
| Male-based | 1R21HD092700-01 | Developing an Animal Model to Identify the Role of the Sperm Centriole in Fertility |
| Female-based | 1R61HD099742-01 | Developing Modulators of the Sperm-specific Potassium Channel Slo3 for Contraception |
| Female-based | 1R43HD094454-01A1 | Development of a Biologic for Non-hormonal Contraception |
| Male-based | 1R21HD083616-01A1 | Development of a Zebrafish Model for Selenoprotein Synthesis and Function |
| Male-based | 1R61HD099720-01 | Development of Allosteric Hipk4 Inhibitors as Non-hormonal Male Contraceptives |
| Female-based | 1R21HD092739-01 | Development of New Therapeutic Strategies for Endometriosis |
| Not-Specified | 1R56HD095629-01 | Development of Novel Sperm-binding Antibodies |
| Not-Specified | 1R01HD084380-01A1 | Developmental Epidemiological Study of Children Born Through Reproductive Technology (Descrt) |
| Not-Specified | 1R21ES024854-01A1 | Dioxin and Estradiol Regulation of Proteins Through Cugbp2 |
| Not-Specified | 1R01GM121688-01 | Dissecting the Origins of a Complex Reproductive Trait: Nematode Self Fertility |
| Male-based | 1R43HD097820-01 | Drug Interactions in Vitro for Ep055: a Non-hormonal Male Contraceptive |
| Female-based | 1R01HD091848-01A1 | Dynamic Regulation of the Ovarian Reserve |
| Male-based | 1F32GM130006-01 | Elucidating the Role of Small Rna Pathways in Heat-stress Induced DNA Damage During Spermatogenesis |
| Male-based | 1F30HD097961-01 | Elucidating the Role of Tcf21+ Mesenchymal Cells in Testis Tissue Homeostasis and Regeneration |
| Male-based | 1R21ES026778-01A1 | Embryonic Inheritance of Sperm Methylome After Adult Exposure to Phthalates |
| Female-based | 1R01ES026998-01A1 | Environmentally Relevant Phthalate Exposures and Ovarian Function |
| Male-based | 1R01GM122776-01A1 | Epigenetic Regulation of Gene Expression During Spermatogenesis |
| Male-based | 1F32HD086939-01A1 | Epigenetic Regulation of Histone Eviction in Spermatogenesis |
| Female-based | 1F32HD095618-01 | Epithelial Regeneration in the Adult Oviduct |
| Male-based | 1F32HD097932-01A1 | Examining the Androgenic and Progestational Effects of Novel Androgens for Male Contraception |
| Male-based | 1R01HD094698-01A1 | Exploring Vascular-mesenchymal Interactions in the Stem Cell Niche |
| Not-Specified | 1R01HD089932-01A1 | Fa DDR Pathway in Germ Line Integrity |
| Male-based | 1R21HD094322-01 | Feasibility of in-home Semen Testing in a North American Preconception Cohort Study |
| Female-based | 1R03HD097262-01 | Fertility Preservation in Women With Cystic Fibrosis Pre-lung Transplantation |
| Female-based | 1R56HD086054-01A1 | FSH and IGF1R Signaling Crosstalk in Ovarian GCS |
| Male-based | 1R15GM126458-01 | Functional Analysis of MicroRNAs in C.elegans Spermatogenesis |
| Male-based | 1R01HD088412-01 | Functional Analysis of Novel Testis-expressed Secreted and Transmembrane Proteins |
| Female-based | 1R01ES029464-01A1 | Functional and Epigenetic Effects of Preconceptional EDCs on the Female HPG Axis |
| Not-Specified | 1P01HD087157-01A1 | Functional Genomics and Dec-tec to Identify Germ Cell-specific Contraceptives |
| Male-based | 1R01HD095341-01 | Functional Genomics and Proteomics to Reveal Reproductive-tract Specific Proteins |
| Not-Specified | 1R01GM117155-01 | Functions of Polo-like Kinases During Mammalian Meiosis |
| Male-based | 1R01ES030722-01A1 | Genetic and Epigenetic Mechanisms of Infertility Caused by Endocrine Disrupting Chemicals |
| Female-based | 1R01HD086175-01A1 | Genetic Interrogation of Conserved Follicular Factors for Matrix Metalloproteinase Regulation and Ovulation |
| Female-based | 1R21HD087427-01A1 | Genetic Studies of the Kit Receptor Tyrosine Kinase in Primordial Follicle Activation |
| Female-based | 1R01HD093778-01 | Genetics and Genomics of the Ovarian Reserve and Female Fertility |
| Female-based | 1R21HD087973-01 | Genome-wide Identification of Genes Required for Decidualization |
| Female-based | 1F31HD088053-01 | Genotoxic Effects of L1 Retrotransposition Trigger Oocyte Elimination During MPI |
| Not-Specified | 1R21AA024889-01A1 | Germ Cell Mediated Epigenetic Memory of Ethanol Exposure |
| Female-based | 4R00HD080742-03 | Greb1 Action in Endometrial Function and Dysfunction |
| Male-based | 1R21ES027117-01 | Histone Lysine Crotonylation in Paternal Epigenetic Inheritance |
| Female-based | 1R01HD093726-01 | Homeostatic to Reactive Hyaluronan Matrices in Ovarian Reproductive Aging |
| Male-based | 1R01ES025066-01A1 | Human Pesticide Exposure and Epigenetic Changes in Sperm DNA |
| Male-based | 1R03HD097433-01 | Identification of Genetic Factors Contributing to Germline Stem Cell Maintenance |
| Male-based | 1R21HD086839-01A1 | Identification of Phospho-proteins Regulating Sperm Function |
| Not-Specified | 1R01ES027921-01A1 | Identity, Mechanisms and Early Life Impacts of Transporter Interfering Compounds |
| Not-Specified | 1F32HD094500-01 | Improving Bovine Cloning Efficiency by Enhancing Reprogramming During Embryonic Genome Activation (EGA) |
| Female-based | 1R01HD089957-01 | Improving Emergency Contraceptive Effectiveness in Obese Women |
| Male-based | 1R01HD092084-01 | Improving Fertility Preservation in Boys With Cancer |
| Not-Specified | 1R01EB027099-01 | In Vivo Analysis of Mammalian Fertilization |
| Female-based | 1R61AI136008-01 | Ind-enabling Preclinical Development of a Multipurpose Intravaginal Ring for the Prevention of Herpes, HIV and Unintended Pregnancy |
| Female-based | 2R44AI124815-02A1 | Ind-enabling Preclinical Development of a System for the Multipurpose Prevention of HIV and Unintended Pregnancy |
| Female-based | 4R00NR017191-02 | Influence of Diet, Iron Stores, and Toxic Metals on Uptakes and Effects on Uterine Fibroid Risk in African American Women |
| Female-based | 1R41HD100190-01 | Innovative 3d Printed Intravaginal Ring Anelleo-pro, the First Single Administration of Progesterone for Infertility |
| Female-based | 1R21HD097601-01 | Integrative Metabolism of Oocyte Development and Its Modulation by Maternal Diet |
| Female-based | 1R21ES026282-01A1 | Investigating Modes of Action of Glyphosate-induced Ovotoxicity |
| Male-based | 1F31GM117971-01 | Investigating the Functional Role of Zinc Fluxes During Sperm Activation |
| Female-based | 5F31HD096838-01 | Investigating the Role of an FSHB Enhancer in FSHB Expression and Polycystic Ovary Syndrome |
| Male-based | 3R01HD091068-03S1 | IPSC-derived Organoids to Study Testis Function |
| Male-based | 1R03HD094046-01 | Isolation of Viable Human Sperm From Failed Microsurgical Testicular Biopsies |
| Male-based | 1R03HD099412-01 | L1 Retrotansposition During Spermatogenic Failure |
| Female-based | 1R01HD093671-01 | Large-scale Studies in Emerge to Discover the Genetic Determinants of Uterine Fibroids |
| Male-based | 1R03HD090315-01 | Lifestyle and Psychosocial Determinants of Male Subfertility: a Prospective Study |
| Male-based | 1R01HD084353-01A1 | Linking Fertility-associated Gene Polymorphisms to Aberrant Sperm Phenotypes |
| Female-based | 1R61AI142687-01 | Long Acting Film Technology for Contraception and HIV Prevention (Latch) |
| Male-based | 1R44HD099040-01 | Low Cost, Automated Smartphone Based Assay for Semen Analysis |
| Male-based | 1R44HD093493-01 | Low-cost, Portable and Automated Semen Analysis Using Computational Microscopy for Home-based Testing of Male Wellness and Fertility |
| Male-based | 1R35GM119458-01 | Macrophage Regulation of the Spermatogonial Stem Cell Niche |
| Male-based | 1R01ES028214-01 | Male Preconception Phthalates and Offspring Embryo and Sperm Allele-specific Methylome Programming |
| Male-based | 1P01HD087157-01A1<br>8121 | Manipulation of Sperm-specific Proteases Using Genetic and Chemical Approaches |
| Female-based | 1R35GM131810-01 | Maternal Control of Germline Development |
| Not-Specified | 1R35GM118092-01 | Mechanism and Regulation of Meiotic Recombination |
| Not-Specified | 3R01DK047320-22S2 | Mechanism of Selenoprotein Synthesis |
| Female-based | 1R01GM124519-01A1 | Mechanisms Driving the Transition From Oocyte to Embryo: the Role of the mRNA Decay Activator Zfp36l2 |
| Female-based | 1F30HD100126-01 | Mechanisms Linking Global Transcriptional Silencing and Zygotic Gene Activation During the Oocyte-to-embryo Transition in Mammals |
| Male-based | 4R00HD081204-02 | Mechanisms of a Novel Actin Related Protein in Male Gametes Ensuring Fertility |
| Not-Specified | 1R01ES027487-01 | Mechanisms of Memory of Environmental Exposure: Determining the Role of Histone Modifications in Regulating Transgenerational Behavior Effects Caused by Environmental Chemical Exposure. |
| Not-Specified | 1R15HD095735-01 | Mechanisms Underlying Telomere Function in Germ Cell Development |
| Female-based | 1R03HD087528-01A1 | Membrane Estrogen Receptor 1 Mediation of Epigenetic Effects of Estrogen |
| Male-based | 1R01HD093827-01A1 | Membrane Lipid Regulation of Calcium Channels in Sperm. |
| Female-based | 1R01HD087402-01A1 | Metabolic Events Controlling Ovarian Steroidogenesis |
| Female-based | 1R01HD092263-01 | Metabolic Regulators of Corpus Luteum Function |
| Male-based | 1R01GM132490-01 | Mitochondria-anchored Protein Complexes in piRNA Biogenesis and Function |
| Female-based | 1F32HD095550-01 | Modelling Human Embryonic Aneuploidy Using Pre-implantation Bovine Embryos |
| Female-based | 1F30AG058387-01A1 | Modelling Ovarian Ageing Phenotype in Mechanically Tuned 3d Matrices |
| Female-based | 5F30DK108561-02 | Molecular and Cellular Changes in the Peritoneal Surface Mesothelium During Adhesion Formation |
| Female-based | 1R21HD094096-01 | Molecular Identity of Maternal Regulators of the Egg to Embryo Transition |
| Male-based | 1R03HD098314-01A1 | Molecular Marker for Centriole Remodelling in Human Reproduction |
| Female-based | 1R35GM122580-01 | Molecular Mechanisms of the Maternal to Zygotic Transition |
| Female-based | 1R43HD097720-01 | Multi-center Prospective Study Assessing a Panel of MicroRNAs as a Non-invasive Test for Endometriosis |
| Female-based | 1R01HD086100-01A1 | Neuroendocrine Regulation of Reproduction by Glucocorticoids |
| Female-based | 1R01HD097675-01A1 | Neurotensin: a Novel Mediator of Ovulation |
| Female-based | 1R61HD099747-01 | Nonhormonal Contraceptive Intravaginal Ring Based on High Valency Anti-Sperm Antibody Constructs |
| Female-based | 1R43HD097941-01 | Non-hormonal Contraception Based on Vaginal Delivery of Multimeric Sperm-binding Antibodies |
| Female-based | 1R03HD098441-01 | Novel Reverse Genetics Approach to Probe Cytoskeletal Functions in Mammalian Oocytes |
| Male-based | 1R01HD090007-01 | Origin and Functional Significance of the Spermatogonial Stem Cell Transcriptome Barcode |
| Female-based | 1R21HD094983-01 | Oxygen as a Potential Regulator of Follicle Quiescence in the Primate Ovary |
| Female-based | 1R03HD090624-01A1 | P21-activated Kinase as Regulator of Actin and Microtubules in Mammalian Oocytes |
| Female-based | 6R21HD080148-02 | Parous Mouse: a Unique Model to Define Uterine Receptivity Versus Nonreceptivity |
| Male-based | 1R01ES028298-01A1 | Paternal Preconception Phthalates and Reproductive Health - Potential Mediation Through Sperm DNA Methylation |
| Female-based | 1F32HD097939-01 | Paxillin as a Mediator of Non-classical Androgen Receptor Actions in Ovarian Follicles |
| Not-Specified | 5R01HD086478-03 | Phenotyping Early Embryonic Lethal Knockout Mice to Identify Essential Genes With Previously Uncharacterized Roles in Pre-implantation Development, Gastrulation, Turning, and Placentation |
| Female-based | 1K99ES028748-01 | Phthalate-induced Ovulatory Dysfunction in Women |
| Female-based | 4R00ES028748-03 | Phthalate-induced Ovulatory Dysfunction in Women |
| Female-based | 1R56ES025147-01A1 | Phthalates and Ovarian Toxicity |
| Female-based | 1R01ES028661-01A1 | Phthalates and Ovarian Toxicity |
| Female-based | 1R56ES025728-01A1 | Placental Molecule Secretions Measured in Early Pregnancy Are Targets of Endocrine Disruption and Are Indicators of Sex-specific Fetal Development |
| Female-based | 1R43HD097944-01 | Plant-derived Molecular Condom as an on-demand Nonhormonal Female Contraceptive |
| Not-Specified | 1R01HD083260-01A1 | Postnatal Plasticity in the GNRH System |
| Female-based | 1R43AI124815-01 | Preclinical Development of a Multipurpose Intravaginal Ring for the Prevention of HIV and Unintended Pregnancy |
| Female-based | 1R01HD100336-01 | Pre-IVF Treatment With a GNRH Antagonist in Women With Endometriosis - a Prospective Double Blind Placebo Controlled Trial (Pregnant) - Application 1/4 |
| Female-based | 1R01HD100329-01 | Pre-IVF Treatment With a GNRH Antagonist in Women With Endometriosis - a Prospective Double Blind Placebo Controlled Trial (Pregnant) - Application 4/4 |
| Female-based | 1R01HD100343-01 | Pre-IVF Treatment With a GNRH Antagonist in Women With Endometriosis - a Prospective Double Blind Placebo Controlled Trial 2/3 |
| Female-based | 1R01HD099487-01 | Primary Ovarian Insufficiency: Etiology and Comorbid Disease |
| Male-based | 1R01HD095631-01 | Project One: Development and Testing of Hca lvr |
| Male-based | 5R01HD095630-02 | Project Three: Assessing Effects of Anti-CD52G MABS on STD Pathogens in Semen |
| Female-based | 1R21HD091337-01 | Prospective Randomized Trial of Tranexamic Acid Versus Levonorgestrel Intrauterine System for the Treatment of Heavy Menstrual Bleeding in Women With Uterine Fibroids |
| Female-based | 1R01HD092499-01A1 | Regulation of Ca<sup>2+</sup> Influx in Mouse Oocytes and Eggs During Maturation and Fertilization to Improve Assisted Reproductive Technologies and Modulate Fertility |
| Not-Specified | 1R01GM123556-01A1 | Regulation of Developmental Potency by the Transposon Line1 |
| Male-based | 1R35GM118052-01 | Regulation of Meiosis in Mice |
| Female-based | 1R15HD099859-01 | Regulation of Oocyte Development by Steroid Hormones |
| Not-Specified | 1R01GM113001-01A1 | Regulation of Sex-specific Gonad Stem Cell Niche Development by Doublesex |
| Male-based | 1R03HD096176-01 | Regulation of Sperm Metabolism and Fertility by Calcineurin and Gsk3 |
| Male-based | 1F31HD097928-01A1 | Replication-independent DNA Methylation Dynamics During Post-Testicular Sperm Maturation |
| Male-based | 1R01GM124024-01 | Requirements for Cytosolic Chaperones in the De Novo Folding of Septin Proteins |
| Male-based | 4R00ES025231-03 | Retinoic Acid Signaling Disruption by Phthalates in Human and Rodent Fetal Testis |
| Male-based | 1K99ES025231-01A1 | Retinoic Acid Signaling Disruption by Phthalates in Human and Rodent Fetal Testis |
| Female-based | 1R01HD098200-01 | Reversible Contraception by Selective Silencing of GNRH-II |
| Female-based | 1R01HD084478-01A1 | Risk Factors for Early Pregnancy Loss |
| Male-based | 1R01HD094546-01A1 | Rna Pol II Pausing Is Critical for Spermatogenesis and Male Fertility |
| Not-Specified | 1R01GM125800-01 | Role of Chromosomally Tethered Proteasome in Chromosome Pairing and Meiotic Recombination |
| Not-Specified | 1R01GM127569-01A1 | Role of GCNA in Preserving Genome Integrity and Fertility |
| Female-based | 1R01HD089495-01A1 | Role of Neuroestradiol in Regulation of the GNRH Surge |
| Female-based | 1R56HD093383-01A1 | Role of the DNA Helicase LSH in Female Meiosis |
| Male-based | 1R01HD093703-01A1 | Role of the Extracellular Matrix During Wolffian/epididymal Duct Morphogenesis |
| Male-based | 1R01HD094736-01A1 | Roles of X- and Y-palindromic Genes in Mammalian Fertility |
| Female-based | 1R44HD095724-01 | Safety, and Acceptability of a Non-hormonal Vaginal Ring |
| Female-based | 1R01HD097202-01A1 | Salt-inducible Kinase Regulation of Ovarian Granulosa Cells |
| Female-based | 1R01GM129478-01 | Sexual Identity Maintenance in Drosophila Female Germ Cells |
| Female-based | 1R44HD097750-01 | Sirt1 and Bcl6: Dual Biomarkers of Endometriosis and Endometrial Receptivity |
| Not-Specified | 1R01HD095628-01 | Slo3 K Channel: a Novel Target for Contraception |
| Female-based | 1R61HD099748-01 | Smart Polymer Fibers for Tampon-like Nonsteroidal Contraceptive Devices |
| Not-Specified | 1R15GM117548-01 | Specification of Meiotic Cohesin Function by Divergent Alpha-kleisin Subunits |
| Male-based | 1R44HD095355-01 | Sperm Sample Preparation for Point of Care Applications |
| Female-based | 1R21HD092009-01 | Stem Cell-derived GNRH Neurons: Optimization and Characterization |
| Female-based | 1R01HD088638-01 | Study of Ovarian Aging and Reserve in Young Women (Soar) |
| Female-based | 1R61HD099745-01 | Synthetic mRNA-mediated Reversible Immunocontraception |
| Not-Specified | 1K99ES025280-01A1 | Systems Approach to Define AHR Ligand Toxicity on Reproductive System Development |
| Male-based | 1P01HD087157-01A1<br>8119 | Targeting Sperm-specific Proteins During Meiosis and Sperm Morphogenesis |
| Male-based | 1R61HD099722-01 | Targeting Testis-specific Ubiquitin-proteasome Pathways for Male Contraception |
| Female-based | 1R01HL134840-01 | Telomeres and Fecundity |
| Female-based | 1R01HD097087-01 | The Actions of Steroid Hormones on Oviduct Function |
| Male-based | 1R01ES028712-01 | The Effects of Environmental Exposures on Semen Quality and the Sperm Epigenome |
| Female-based | 1R03HD095098-01 | The Function of Progesterone Receptor in Human Ovulatory Follicles |
| Female-based | 1F32HD098805-01 | The Homeodomain Transcription Factors, Six6 and Six3, in the Circadian Regulation of Reproduction |
| Not-Specified | 7R21HD088792-02 | the Impact of Piwi Associated Transcripts in Xenopus Germ Cell Development |
| Female-based | 1R15HD087911-01 | The Interaction Between Nat2 Acetylator Status and Exposure to Tobacco Smoke on Ovarian Reserve and in Vitro Fertilization Outcomes. |
| Female-based | 1R01HD091117-01A1 | The Oocyte's Progression Through Meiosis: Involvement of a Heart Disease-associated Protein |
| Female-based | 1R01HD096077-01A1 | The Role of FOS in the Ovary |
| Male-based | 1R03HD090306-01 | the Role of Transcription Factor S-sox5 in Male Fertility and Sperm Flagella Formation |
| Female-based | 1R21HD094293-01A1 | Timing Endometrial Receptivity |
| Female-based | 1R01HD098278-01 | Towards a Preclinical Model for Overcoming Infertility With Induced Pluripotent Stem Cells |
| Female-based | 3R01ES027051-02S1 | Transdisciplinary Approach to Rapidly Identify Reproductive Toxics in Pregnant Women and Newborns |
| Male-based | 1R03OH011540-01 | Transgenerational Work Exposures, EDCs and Male Fertility |
| Male-based | 1R44HD100256-01 | Translational Feasibility of an Oral, Non-hormonal, Male Contraceptive Pill |
| Male-based | 1R56AG052581-01A1 | Tumor Suppressors Mediate a Reduction in Male Gamete Quality With Aging |
| Female-based | 1R21ES028963-01A1 | Water Disinfection by-products and Female Fertility |
| Male-based | 1R01GM130691-01 | Y Chromosome Evolution |

**Supplementary Table VIII.** The NIHR’s Total Funding Awarded for the 21 health categories of research and specialties as of April 1*^st^*, 2011. The present data is extracted from the NIHR’s Open Data.

| **HRCS Health Categories** | **Total Funding Awarded (£)** |
| --- | --- |
| **Mental Health** | 493,552,549 |
| **Generic Health Relevance** | 484,950,225 |
| **Cancer and Neoplasms** | 347,947,217 |
| **Infection** | 275,265,348 |
| **Cardiovascular** | 224,898,509 |
| **Neurological** | 222,002,645 |
| **Reproductive Health and Childbirth** | 216,618,019 |
| **Oral and Gastrointestinal** | 164,146,703 |
| **Respiratory** | 146,665,215 |
| **Metabolic and Endocrine** | 129,821,105 |
| **Musculoskeletal** | 126,596,631 |
| **Stroke** | 121,450,341 |
| **Injuries and Accidents** | 119,415,772 |
| **Renal and Urogenital** | 108,827,444 |
| **Skin** | 66,127,071 |
| **Eye** | 63,433,235 |
| **Inflammatory and Immune System** | 54,122,033 |
| **Blood** | 29,836,676 |
| **Disputed etiology and other** | 23,847,721 |
| **Congenital Disorders** | 18,740,901 |
| **Ear** | 11,238,128 |

**Supplementary Table IX.** The NIH fund 299 various research, conditions, and diseases categories (RCDC) and a project can fall under multiple RCDCs as the NIH does not budget research per category NIH, 2021). The estimated proportion of funding for the chosen categories of contraception/ reproduction, infertility, obesity, and mental health against the annual NIH total research grant funding between FY2016 to FY2019. The FY2020 was excluded from this table as only 2 projects were collected and would be incomparable. Value ($000) refers to the estimated funding by the NIH RCDC Categorical Spending in millions. (%) is the calculated funding percentage or proportion for the RCDCs from the annual NIH Total Research Funding

| **Supplementary Table IX : Estimated Proportion of Funding to RCDC by the NIH against the NIH's Total Budget** |  |  |  |  |  |
| --- | --- | --- | --- | --- | --- |
| **RCDC** |  | **Fiscal Years** |  |  |  |
|  |  | **2016** | **2017** | **2018** | **2019** |
| **Contraception/**<br>**Reproduction** | Value (\$000) | 419,000 | 437,000 | 496,000 | 547,000 |
|  | (%) | 1.85 | 1.82 | 1.91 | 1.94 |
| **Infertility** | Value (\$000) | 86,000 | 94,000 | 120,000 | 151,000 |
|  | (%) | 0.38 | 0.39 | 0.46 | 0.54 |
| **Obesity** | Value (\$000) | 965,000 | 999,000 | 1,055,000 | 1,109,000 |
|  | (%) | 4.26 | 4.16 | 4.07 | 3.94 |
| **Mental Health** | Value (\$000) | 2,454,000 | 2,717,000 | 3,010,000 | 3,296,000 |
|  | (%) | 10.83 | 11.31 | 11.62 | 11.71 |
| **Total NIH Research Funding (\$)** | | 22,649,752,290 | 24,031,670,764 | 25,906,788,735 | 28,143,252,479 |

**Supplementary Figure 1.**
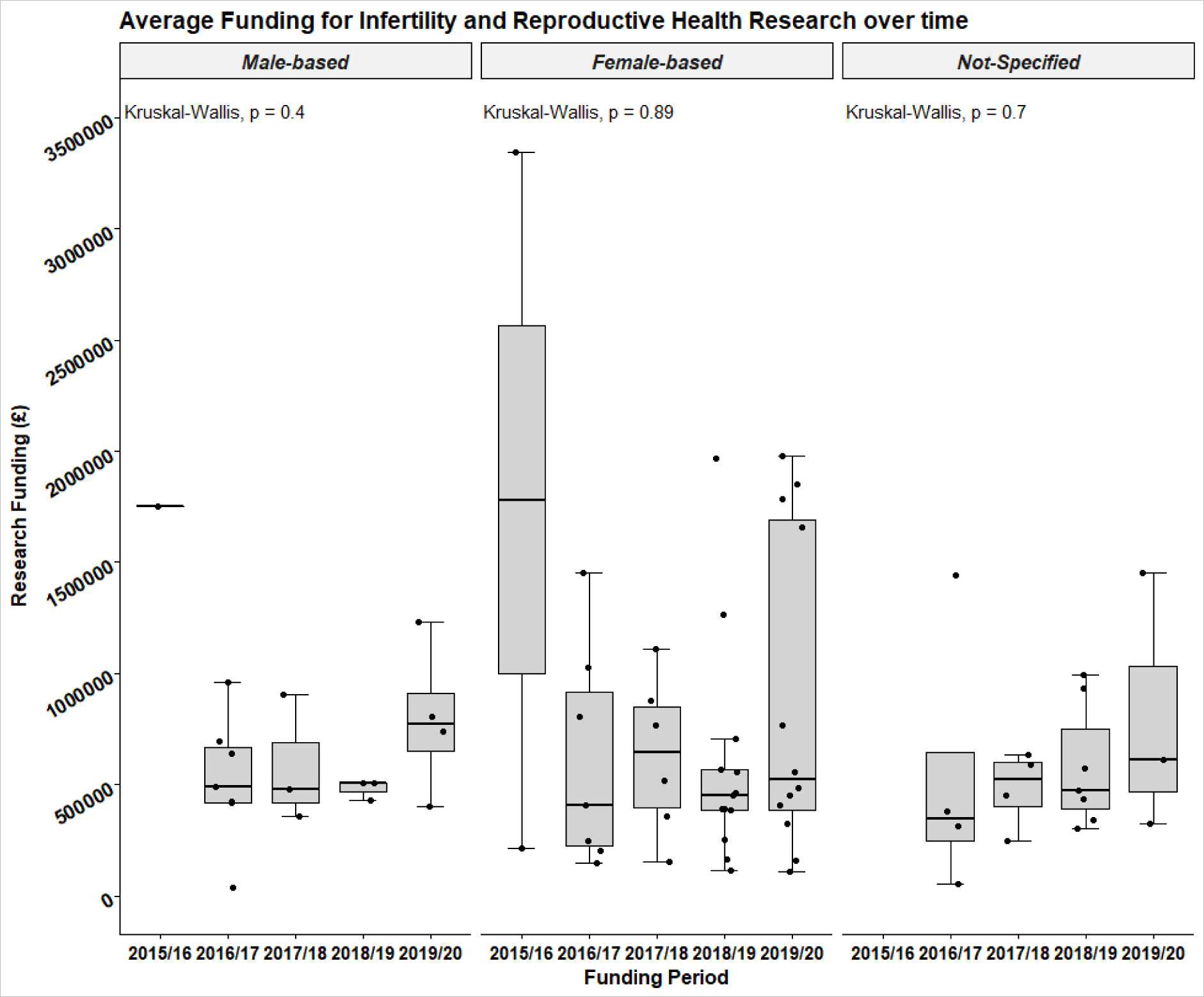
The UK average funding across consecutive funding periods box-and-whisker plot with a 95% CI. 3 projects were collected between January 1^st^ to March 31^st^, 2016, for the 2015/16 funding period, therefore many projects were not expected to be awarded funding. No statistically significant differences of funding variation were observed by the Kruskal-Wallis test over the consecutive funding periods of each research group. In the male-based group, P=0.39 and χ^2^=4.08 with 4 degrees of freedom. For the female-based group, P=0.89 and χ^2^=1.1 with 4 degrees of freedom. In the not-specified group, P=0.7 and χ^2^=1.41 with 3 degrees of freedom.

**Supplementary Figure 2.**
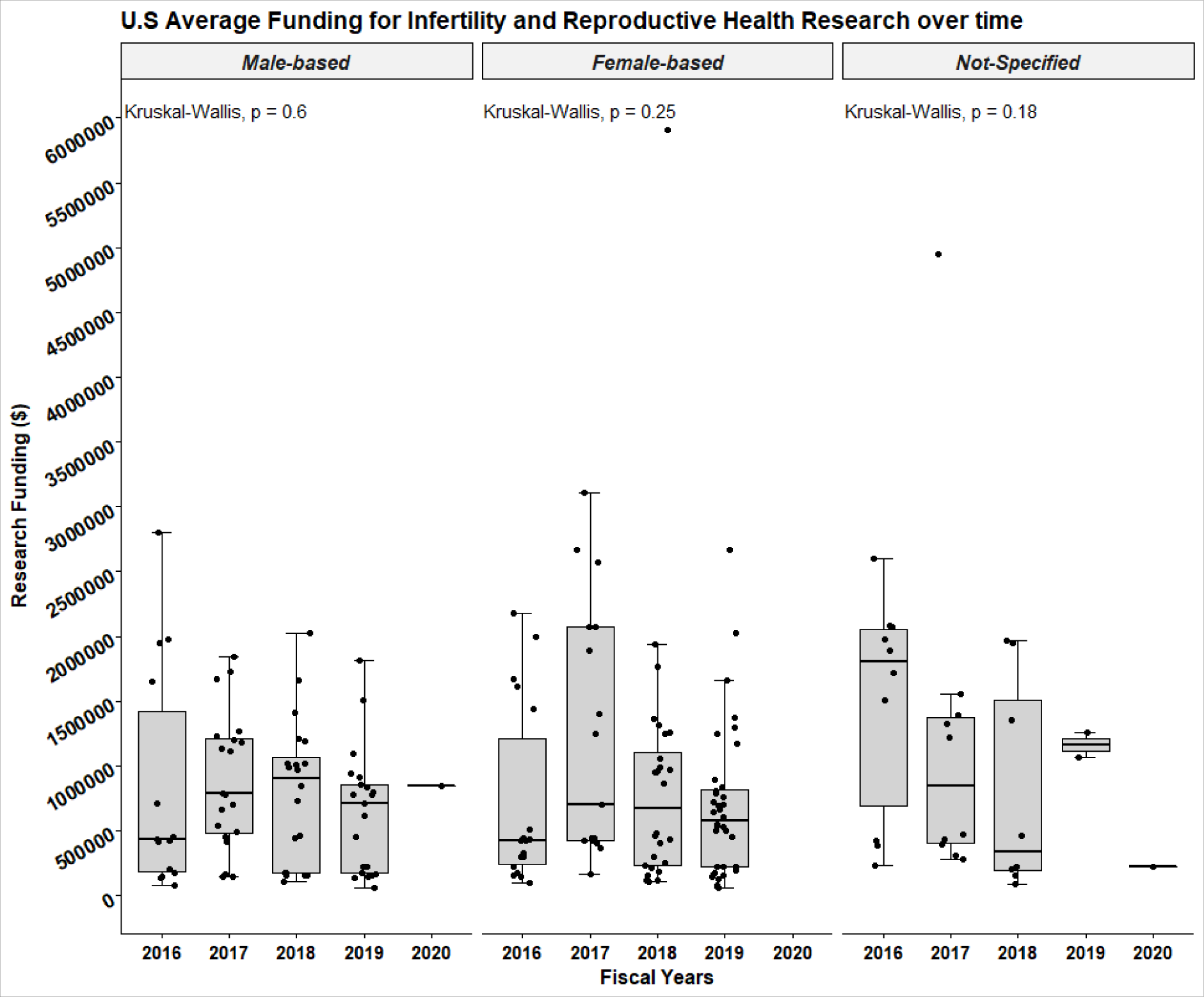
Box-and-whisker plot has a 95% CI for the funding awarded across the five consecutive FYs by the NIH institutes. In the FY2020, 2 projects were awarded funding between 1^st^ October to 31^st^ December 2019. The Kruskal-Wallis did not observe statistically significant differences of funding variation over the consecutive FYs of each research group. For the male-based group, P=0.59 and χ^2^=2.76 with 4 degrees of freedom. In the female-based group, P=0.25 and χ^2^=4.12 with 3 degrees of freedom. In the not-specified group, P=0.18 and χ^2^=6.28 with 4 degrees of freedom.

## References

1. Barratt CLR, Bjorndahl L, De Jonge CJ, Lamb DJ, Osorio Martini F, McLachlan R, Oates RD, van der Poel S, St John B, Sigman M, Sokol R, Tournaye H. The diagnosis of male infertility: an analysis of the evidence to support the development of global WHO guidance-challenges and future research opportunities. Hum Reprod Update 2017; 23: 660–680.

2. Barratt CLR, De Jonge CJ, Sharpe RM. ’Man Up’: the importance and strategy for placing male reproductive health centre stage in the political and research agenda. Hum Reprod 2018; 33: 541–545

3. Barratt CLR, De Jonge CJ, Anderson RA, Eisenberg ML, Garrido N, Rautakallio Hokkanen S, Krausz C, Kimmins S, O’Bryan MK, Pacey AA. A global approach to addressing the policy, research and social challenges of male reproductive health. Human Reproduction Open; Volume 2021, Issue 1

4. Boivin J, Bunting L, Collins JA, Nygren KG. International estimates of infertility prevalence and treatment-seeking: potential need and demand for infertility medical care. Hum Reprod 2007; 22: 1506–1512.

5. De Jonge C, Barratt CLR. The present crisis in male reproductive health: an urgent need for a political, social, and research roadmap. Andrology 2019; 7: 762–76.

6. Liao SJ, Xu YY, Sun RJ, Lyu QY. National natural science foundation of China leads the comprehensive development of basic research in the field of male reproductive health in China. Asian J Androl 2020; 22: 547–548/

7. National Institute for Health and Clinical Excellence. Fertility: Assessment and Treatment for People with Fertility Problems NICE Clinical Guideline, Manchester.2013

8. NIH. National Institutes of Health – Institutes, Centers and Office. [Internet]. 2021. Available from: https://www.nih.gov/institutes-nih/list-nih-institutes-centers-offices

9. NIHR. National Institute for Health Research - About Us.[Internet]. 2021; Available from: https://www.nihr.ac.uk/about-us/

10. NIHR. National Institute for Health Research - Reproductive Health and Childbirth.[Internet]. 2021; Available from: https://www.nihr.ac.uk/explore-nihr/specialties/reproductive-health.htm

11. RePORT – Research Portfolio Online Reporting Tools. Budget and Spending. [Internet]. 2021. Available from: https://report.nih.gov/funding/nih-budget-and-spending-data-past-fiscal-years/budget-and-spending

12. Schlegel PN, Sigman M, Collura B, De Jonge CJ, Eisenberg ML, Lamb DJ, Mulhall JP, Niederberger C, Sandlow JI, Sokol RZ, Spandorfer SD, Tanrikut C, Treadwell JR, Oristaglio JT, Zini A. Diagnosis and treatment of infertility in men: AUA/ASRM guideline part I. Fertil Steril 2021a; 115: 54–61.

13. Schlegel PN, Sigman M, Collura B, De Jonge C, Eisenberg ML, Lamb DJ, Mulhall JP, Niederberger C, Sandlow JI, Sokol R, Spandorfer SD, Tanrikut C, Treadwell JR, Oristaglio JT, Zini A. Diagnosis and treatment of infertility in men: AUA/ASRM guideline part II. Fertil Steril 2021b; 115: 62–69.

14. UKRI. UK Research and Innovation - Annual Report and Accounts 2018–2019. In Department for Business EaIS (ed). 2019, pp. 144

15. [dataset]* Gumerova, Eva; De Jonge, Christopher; Barratt, Christopher (2021), Research Funding for Male Reproductive Health and Infertility in the UK and USA [2016 – 2019], Dryad, Dataset, https://doi.org/10.5061/dryad.v9s4mw6wc

